# Function and contribution of two putative *Enterococcus faecalis* glycosaminoglycan degrading enzymes to bacteremia and catheter-associated urinary tract infection

**DOI:** 10.1101/2024.05.08.593205

**Authors:** Alexandra O. Johnson, Braden M. Shipman, Benjamin C. Hunt, Brian S. Learman, Aimee L. Brauer, Serena P. Zhou, Rachael Hageman Blair, Nicole J. De Nisco, Chelsie E. Armbruster

## Abstract

*Enterococcus faecalis* is a common cause of healthcare acquired bloodstream infections and catheter associated urinary tract infections (CAUTI) in both adults and children. Treatment of *E. faecalis* infection is frequently complicated by multi-drug resistance. Based on protein homology, *E. faecalis* encodes two putative hyaluronidases, EF3023 (HylA) and EF0818 (HylB). In other Gram-positive pathogens, hyaluronidases have been shown to contribute to tissue damage and immune evasion, but function in *E. faecalis* has yet to be explored. Here, we show that both *hylA* and *hylB* contribute to *E. faecalis* pathogenesis. In a CAUTI model, Δ*hylA* exhibited defects in bladder colonization and dissemination to the bloodstream, and Δ*hylB* exhibited a defect in kidney colonization. Furthermore, a Δ*hylA*Δ*hylB* double mutant exhibited a severe colonization defect in a model of bacteremia while the single mutants colonized to a similar level as the wild-type strain, suggesting potential functional redundancy within the bloodstream. We next examined enzymatic activity, and demonstrate that HylB is capable of digesting both HA and CS *in vitro* while HylA exhibits only a very modest activity against heparin. Importantly, HA degradation by HylB provided a modest increase in cell density during stationary phase and also contributed to dampening of LPS-mediated NF-Bκ activation. Overall, these data demonstrate that glycosaminoglycan degradation is important for *E. faecalis* pathogenesis in the urinary tract and during bloodstream infection.

## Introduction

*Enterococcus faecalis* infections are common and impose a high burden on the healthcare system. *E. faecalis* ranks as the 5^th^ most frequently encountered healthcare-associated pathogen in the United States and can cause a variety of infections including endocarditis, central line-associated bloodstream infection (CLABSI), and catheter-associated urinary tract infections (CAUTIs)(1–3). *E. faecalis* is the most common CLABSI pathogen in both pediatric intensive care units and adult long-term acute-care hospitals (1, 2), and is among the three most common causes of CAUTI (3). *E. faecalis* CAUTI can also progress to disseminated infection and secondary bacteremia, and is estimated to cause approximately 13,000 deaths per year in the United States alone (4, 5).

One host factor that may protect against invasive Enterococcal infections are glycosaminoglycans (GAGs). GAGs are acidic polysaccharides that are ubiquitous on the cell surface and extracellular matrix and they provide tissue integrity and structure in addition to mediating immune signaling and tissue repair (6–10). With respect to infection, GAGs are present at many barrier sites such as the endothelial glycocalyx and the urinary tract urothelial lining where they are thought to play a protective role against pathogen invasion (11, 12). For example, the GAG-rich glycocalyx of the bladder mucosa is believed to mask uroplakins and other surface proteins that are targeted by bacteria for adhesion and invasion (11–13). In support of this hypothesis, removal of GAGs from the bladder mucosa increases attachment of *Enterococcus faecalis* as well as other uropathogenic bacteria (14–16).

The most common GAGs in urothelium and endothelial glycocalyx are hyaluronic acid (HA), chondroitin sulfate (CS), and heparin (6, 17). Several pathogenic bacteria produce enzymes that digest GAGs, which can destroy barrier functions, provide a carbohydrate source for microbial growth, and alter the innate immune response either by promoting inflammation or disrupting inflammatory signaling. For example, some *Streptococcus* species produce hyaluronidase enzymes that depolymerize HA and have been shown to promote tissue invasion, disrupt inflammatory signaling, and facilitate bacterial growth by liberating a new carbon source (18–21). We therefore hypothesized that GAG degradation by *E. faecalis* could contribute to pathogenesis and invasive disease.

There are several conflicting reports pertaining to *E. faecalis* hyaluronidase activity. In 1964, Rosan and Williams reported hyaluronidase activity in oral *Enterococcus* isolates (22). However, this study was performed at a time when enterococcal taxonomy was in a state of flux, so it is unclear whether the hyaluronidase-positive strains were *E. faecalis*, *E. faecium*, or another species (23, 24). A putative hyaluronidase was identified in *Enterococcus faecium* that is predominantly associated with vancomycin-resistant clinical isolates (25), and transfer of the plasmid encoding the putative hyaluronidase increased virulence of a fecal isolate in a mouse peritonitis model (26). However, no hyaluronidase activity was observed for any of the *E. faecium* strains (27), and deletion of the putative hyaluronidase itself did not reduce fitness in the mouse peritonitis model (28).

A recent assessment of *in vitro* GAG digestion by several *E. faecalis* clinical isolates found that none of the isolates degraded HA, CS, or HS (29). However, a homology of the *Streptococcus* hyaluronidase gene is present in all 518 *E. faecalis* complete genome sequences available through the Bacterial and Viral Bioinformatics Resource Center (BV-BRC)(30) and 68% of the isolates also have a second putative hyaluronidase. *E. faecalis* strain V587 is one of the clinical isolates with both putative hyaluronidases. This strain is particularly notable as it is vancomycin-resistant and was isolated from the urine of a patient who later experienced sepsis with the same strain (31), making it relevant to two of the most prevalent types of *E. faecalis* infection. The purpose of this study was to characterize the enzymatic activity of both putative hyaluronidases and examine contribution to CAUTI and bacteremia.

## Results

### *Enterococcus faecalis* V587 encodes two putative hyaluronidases with differing predicted protein architectures

The genome of *E. faecalis* strain V587 in the Bacterial and Viral Bioinformatics Resource Center (BV-BRC)(30) contains two putative hyaluronidases located in different parts of the chromosome: *ef3023* (*hylA*) is 4,119 bp at position at 2,884,596 bp (Figure 1A), and *ef0818* (*hylB*) is 3,015 bp at position 763,339 bp (Figure 1B). The *hylA* gene is monocistronic, and no neighboring genes are predicted to be involved in oligosaccharide digestion or uptake. The *hylB* gene is similarly monocistronic but immediately downstream of an operon that encodes for a putative hyaluronate-oligosaccharide phosphotransferase system with a divergently-transcribed GntR-family transcriptional regulator (32).

**FIG 1.**
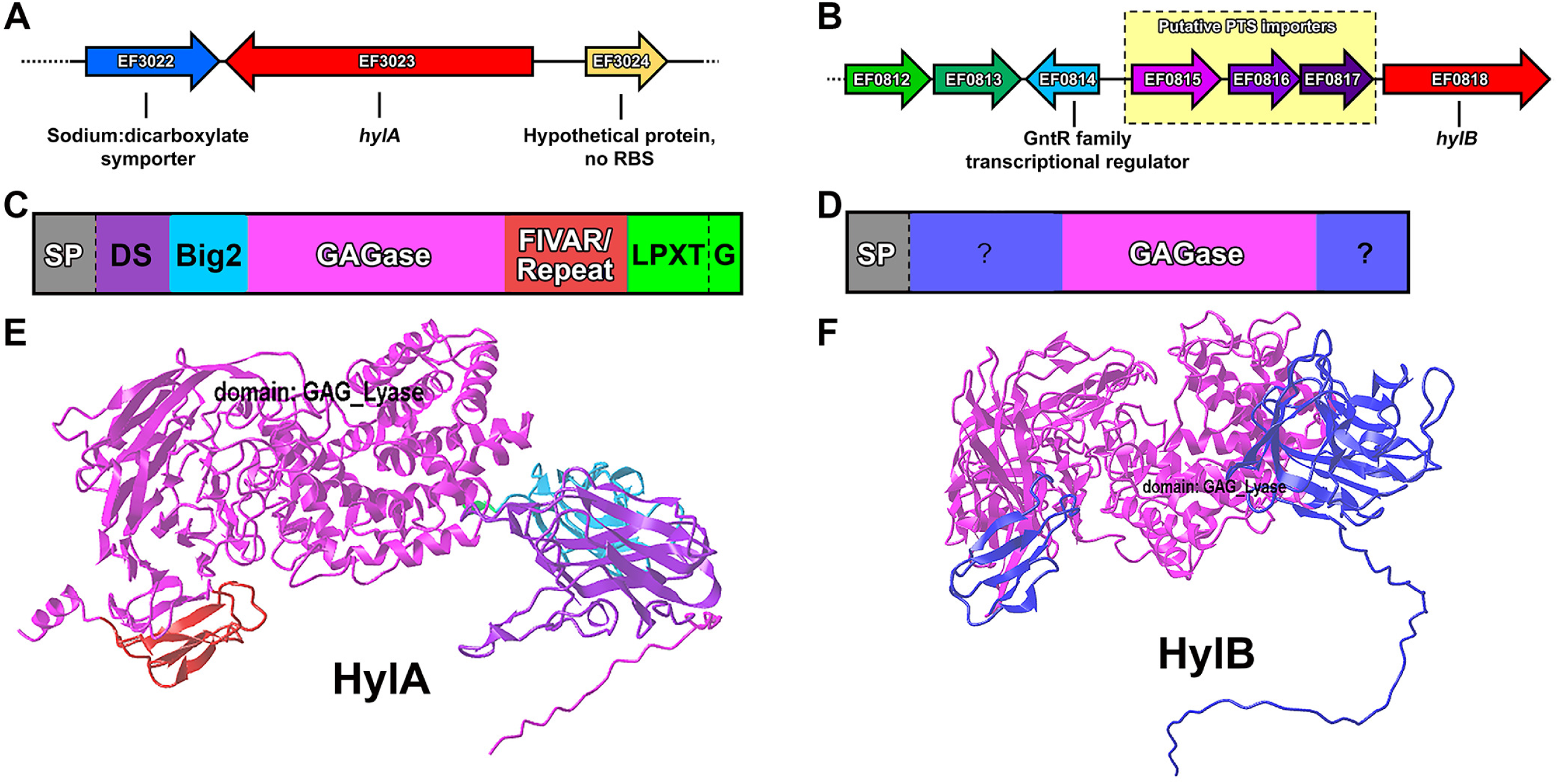
Genetic context and predicted protein architecture of HylA and HylB. Panels A and B show the genetic context of EF3023 (*hylA*) and EF0818 (*hylB*), respectively. Yellow box with dashed outline indicate a putative oligosaccharide phosphotransferase system upstream of *hylB*. Panels C and D show the domain architecture of HylA and HylB, respectively. SP indicates a signal peptide as predicted by SignalP 5.0. A dashed line indicates a predicted proteolytic cleavage site with either signal peptidase (SP) or sortase (LPXTG). The FIVAR/Repeat domain indicates a complete 69 amino acid segment with a predicted “FIVAR” domain, followed by three 71 amino acid repeats with partial FIVAR domain prediction. “?” indicates a stretch of >40 amino acids which have no predicted conserved domains or other obvious features. Panels E and F show AlphaFold structure predictions of HylA and HylB, respectively.

Using the Bacterial and Viral Bioinformatics Resource Center (BV-BRC)(33), we searched 646 complete genome sequences of *E. faecalis* for the presence of *hylA* and *hylB*. A total of 582 strains (90%) had at least one putative hyaluronidase with >95% amino acid identity and >80% query coverage: 470 strains (73%) had a homolog of *hylA*, 459 strains (71%) had a homolog of *hylB,* and 289 (45%) had both. Thus, *hylA* and *hylB* are well conserved across *E. faecalis* strains. Both are predicted to encode “polysaccharide lyase family 8” enzymes, and both are predicted to contain Sec pathway protein export signal peptides according to SignalP 5.0 (34). However, the amino acid sequences of HylA and HylB share only 36% identity to each other and 24-30% identity with hyaluronidases from other bacterial species (Supplemental Table 1). There are also substantial differences in the domain architecture of the two proteins: in addition to the GAGase domain, HylA contains a Discoidin (DS) and Bacterial immunoglobulin-like fold (Big2) domains, a section of repeated amino acid residues annotated as a “Found In Various ARchitectures” (FIVAR) domain, and a predicted LPXTG domain for cell wall attachment (Figure 1C) (35), while HylB lacks cell wall anchoring motifs and has only two small flanking domains with no predicted function by an NCBI Conserved Domain Search (36) (Figure 1D). Structures predictions of HylA (Figure 1E) and HylB (Figure 1F) were generated using AlphaFold (37, 38), with the GAGase domain shown in pink and other domains in different colors. Importantly, an alignment of the two proteins could not be achieved, which underscores the substantial differences in structure.

### *E. faecalis* V587 does not exhibit hyaluronidase activity under *in vitro* conditions

Considering the prior literature regarding a lack of hyaluronidase activity in *E. faecalis* isolates (29), we first sought to examine expression of *hylA* and *hylB* in strain V587. Notably, RNA sequencing data from the closely-related V583 strain indicates that *hylA* and *hylB* are not expressed during growth in BHI (39). We therefore used RT-qPCR to examine expression of *hylA, hylB,* and the housekeeping gene *recA* in V587 after 4 hours of growth in BHI, 0.5X TSB (a condition previously found to support hyaluronidase activity in other species) (29), and human urine (Figure 2). TSB and urine were also supplemented with either unfractionated high molecular weight HA or low molecular weight HA (∼5 kDa) to determine if expression is induced by hyaluronic acid. The Cq values for *recA* were similar across all conditions, confirming that *recA* is an appropriate housekeeping gene for normalizing expression (Figure 2A), and all samples were detected at least 10 cycles before the respective no template controls (Figure 2B). When growth in BHI was used as the calibrator, growth in 0.5X TSB resulted in a slight increase in expression of both *hylA* and *hylB* gene while growth in urine reduced expression (Figure 2C). To examine the impact of HA on expression, we next used unsupplemented 0.5X TSB or urine as the calibrator. In TSB, supplementation with unfractionated HA slightly induced expression of *hylA* but had no effect on *hylB* (Figure 2D). In contrast, unfractionated HA had no impact on expression in urine, but supplementation with 5 kDa HA induced expression of both *hylA* and *hylB* (Figure 2E). Thus, *hylA* and *hylB* are expressed by *E. faecalis* V587 under multiple growth conditions, and expression can be stimulated to some extent by HA.

**FIG 2.**
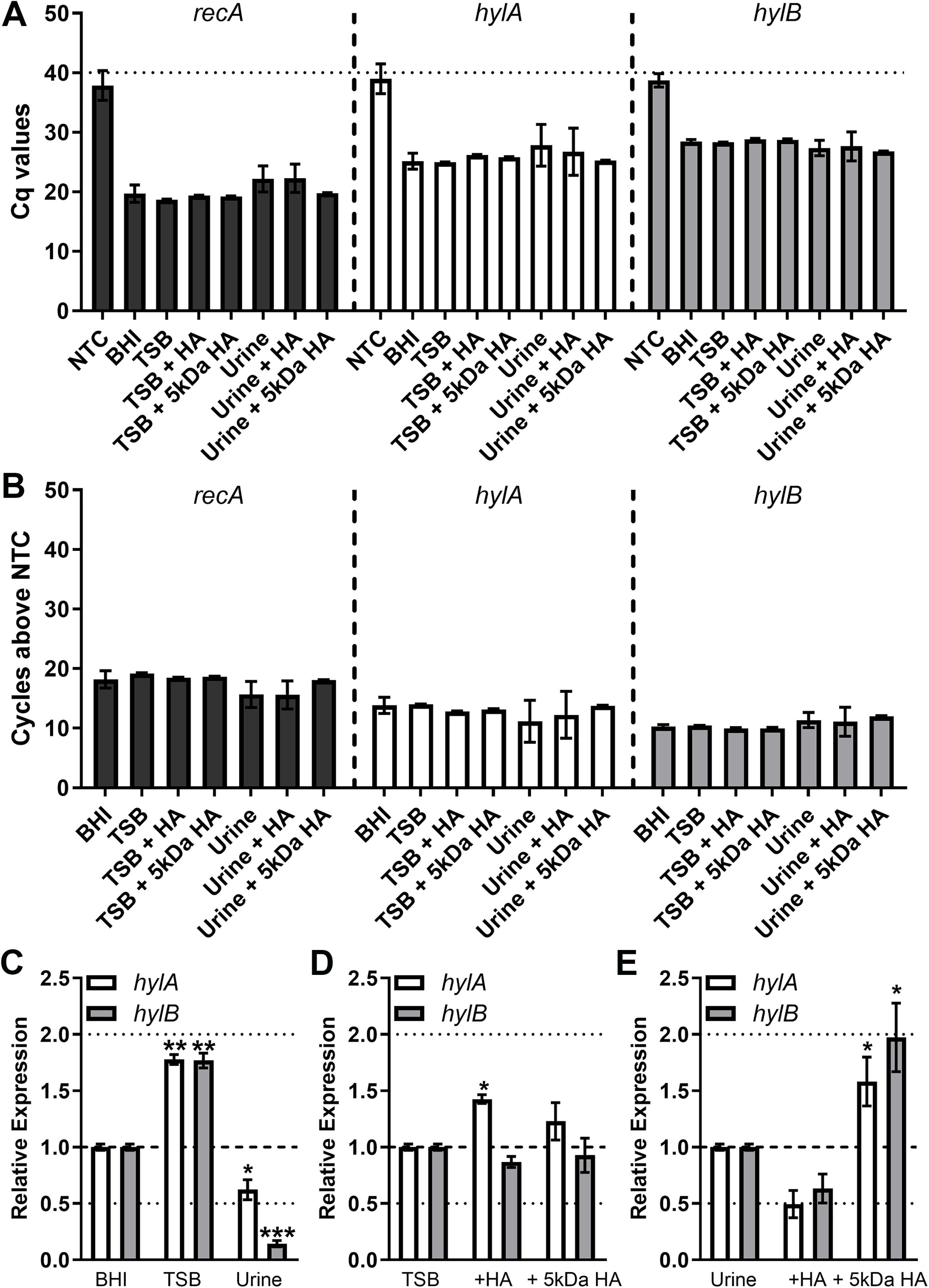
Expression profiles of *hylA* and *hylB* in *E. faecalis* V587. *E. faecalis* strain V587 was cultured in BHI, 0.5X TSB, or human urine with or without supplementation with unfractionated high molecular weight HA or 5 kDa HA, and mRNA levels of *recA, hylA,* and *hylB* were determined by quantitative RT-PCR. (A) Cycle thresholds achieved for each gene compared to a no-template control (NTC). (B) Data display the number of cycles below the NTC at which each transcript was detected. (C) Expression of *hylA* and *hylB* were normalized to the *recA* housekeeping gene and expression across different growth media was compared to expression in BHI by the ΔΔCT method. (D-E) Induction of expression by the presence of HA was determined by comparison to expression in unsupplemented TSB (D) or urine (E). Error bars represent mean ± standard deviation (SD) for at least two independent experiments with two technical replicates each. *\*P*<0.05, *\*\*P*<0.01 by Wilcoxon signed rank test against a hypothetical value of 1.0.

We generated markerless deletion mutants of *hylA* and *hylB*, as well as a double deletion mutant (Δ*hylA*Δ*hylB*). All of the mutants grew similarly to V587 in BHI, TSB, and human urine (Figure 3 A-C). To examine *in vitro* hyaluronidase activity, V587 and the Δ*hylA*Δ*hylB* mutant were incubated on agar plates containing HA and examined for development of a zone of clearance around the colony (Figure 3D). Consistent with prior studies, no hyaluronidase activity was detected for either strain, suggesting that while *hylA* and *hylB* are expressed *in vitro,* active protein is not produced under these conditions. To determine if the lack of activity is specifically due to lack of expression or function under these conditions, *hylB* was cloned into pBAD and transformed into *Lactococcus lactis,* a Gram-positive species that lacks endogenous hyaluronidases. Notably, expression of the V587 *hylB* gene in *L. lactis* resulted in modest hyaluronidase activity, suggesting that the *E. faecalis* HylB can indeed function as a hyaluronidase and the lack of activity in V587 is likely due to regulatory control of expression and activity.

**FIG 3.**
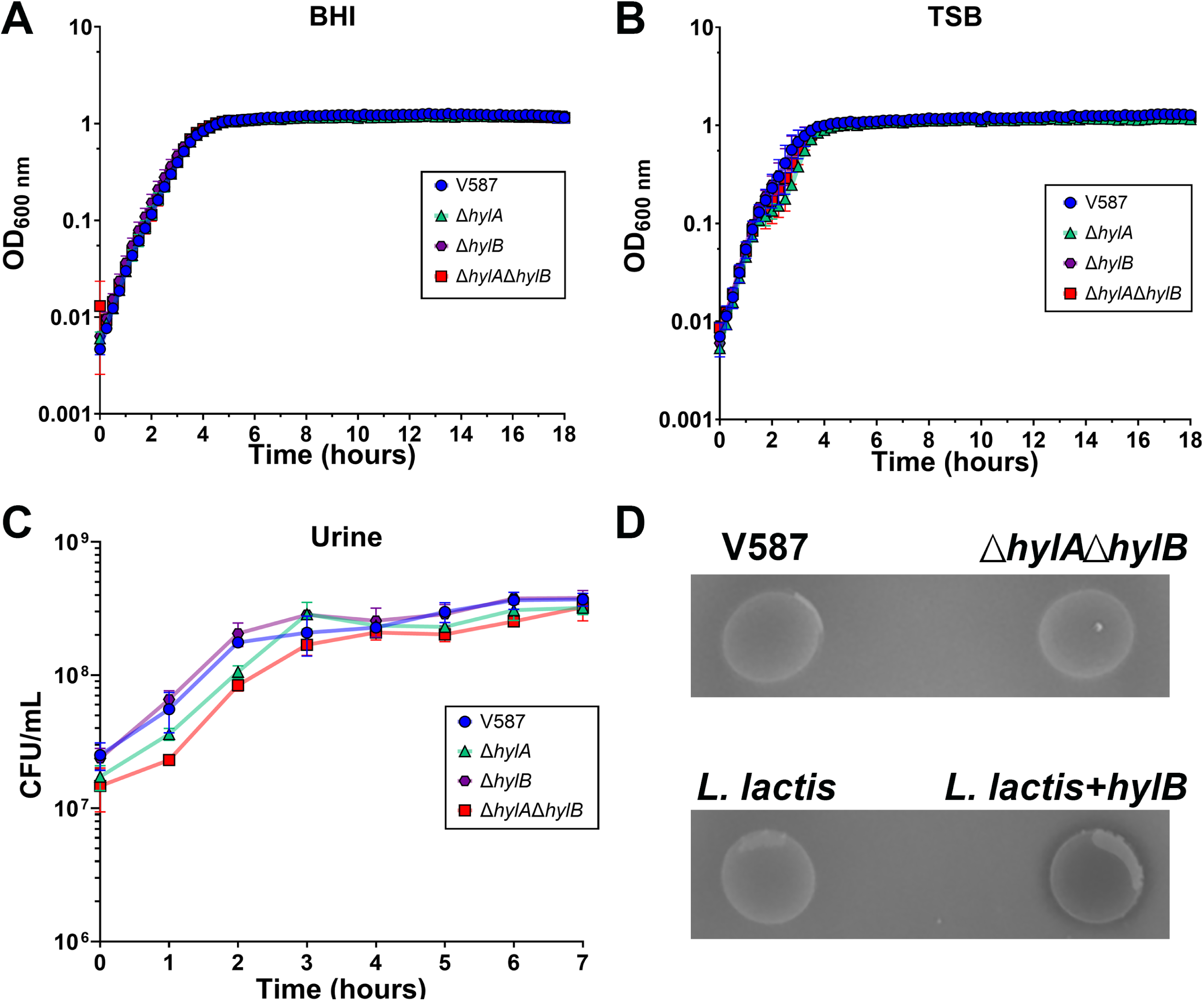
*In vitro* growth rates hyaluronidase activity of *E. faecalis* strains. (A-B) *E. faecalis* V587 and isogenic mutants were grown in BHI broth (A) or TSB (B) at 37°C with shaking and OD600 was measured every 15 minutes for 18 hours. Graphs are representative of two independent experiments. Error bars indicate mean ± SD for three technical replicates. (C) *E. faecalis* V587 and isogenic mutants were grown in human urine at 37°C with shaking and CFUs were determined every hour for 7 hours. Error bars indicate mean ± SD for three independent experiments. (D) Representative images of *E. faecalis* V875, Δ*hylA*Δ*hylB, Lactococcus lactis,* and *L. lactis* constitutively expressing *hylB* after incubation on agar plates containing HA. A zone of clearance is only observed if the bacterium is producing active hyaluronidase.

### HylA and HylB contribute to *E. faecalis* bladder colonization and bacteremia

Prior to further probing the enzymatic activity of HylA and HylB, we first sought determine contribution to *E. faecalis* pathogenesis in our established mouse model of CAUTI. In brief, female CBA/J mice were inoculated transurethrally with 1 x 10^5^ CFU of either V587, Δ*hylA*, Δ*hylB*, or Δ*hylA*Δ*hylB*. During inoculation, a small piece of silicone tubing was left in the bladder to simulate a urinary catheter, as previously described (40). At 48 hours post infection (HPI) the mice were sacrificed and bladders, kidneys and spleens were collected for quantitation of CFUs, with spleen CFU used as an indicator of bacteremia (Figure 4A). Inoculation with the Δ*hylA* mutant significantly reduced bladder and spleen colonization compared to infection with wild-type V587, and also reduced the overall incidence of bacteremia. In contrast, inoculation with the Δ*hylB* mutant resulted in similar bladder colonization as wild-type V587 but a significant defect in the kidneys. Therefore, HylA and HylB both contribute to *E. faecalis* CAUTI but may be differentially expressed in different host niches or have differing roles in pathogenesis. Interestingly, infection with the Δ*hylA*Δ*hylB* double mutant phenocopied the mean bladder colonization of infection with Δ*hylA* and the men kidney colonization as Δ*hylB,* but the only statistically significant defect compared to wild-type V587 was a reduction in spleen colonization and incidence of bacteremia.

**FIG 4.**
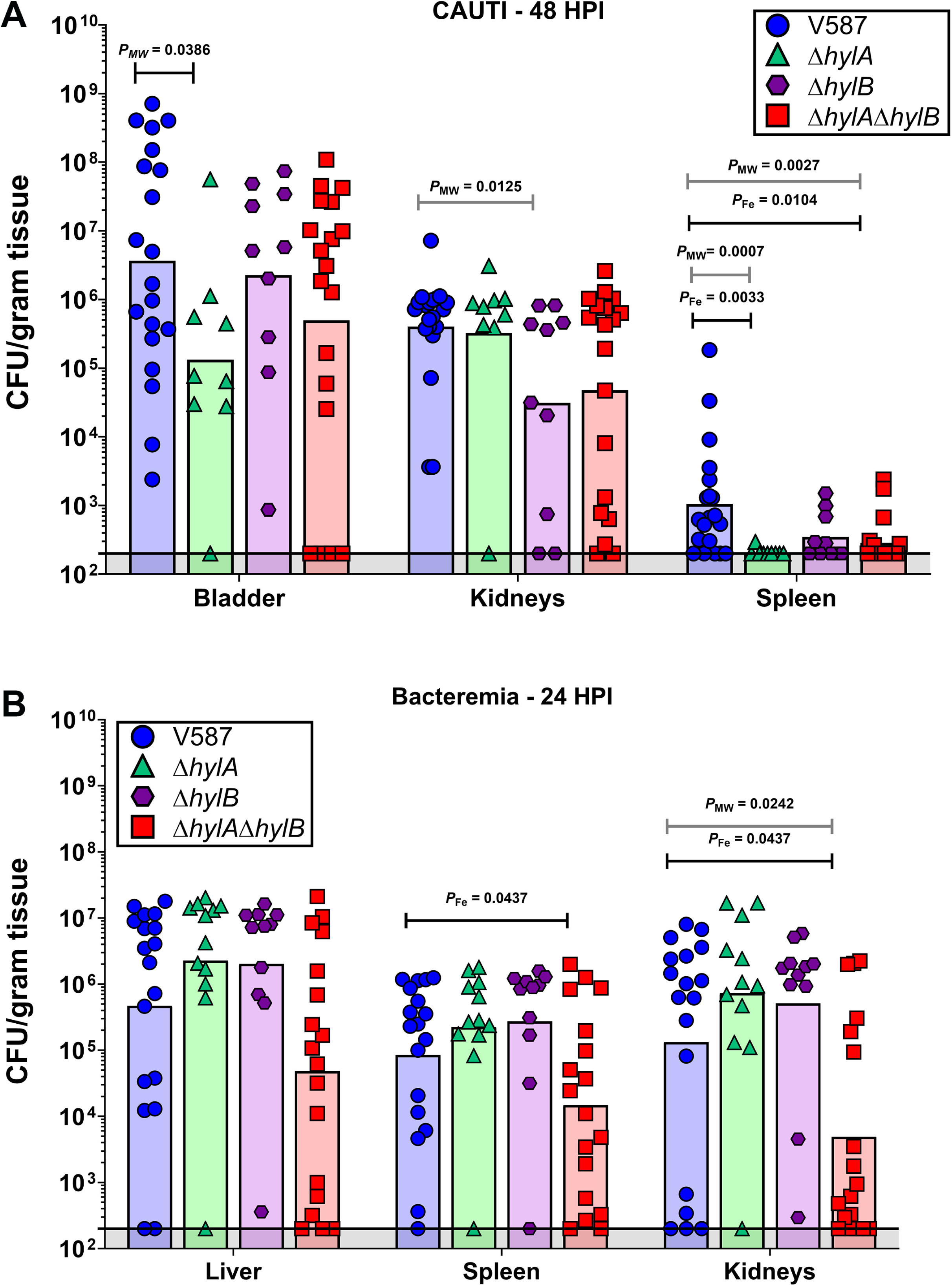
Contribution of *hylA* and *hylB* to CAUTI and bacteremia. (A) 7-week old female CBA/J mice were inoculated via transurethral catheterization with 1 x 10^5^ CFU of V587 (n = 20), Δ*hylA* (n = 9), Δ*hylB* (n = 10), or Δ*hylA*Δ*hylB* (n = 19). CFUs were determined for bladder, kidneys, and spleen 48 HPI. (B) 7-week old female CBA/J mice were inoculated via tail vein injection of 1 x 10^8^ CFU of V587 (n= 18), Δ*hylA* (n = 12), Δ*hylB* (n = 11), or Δ*hylA*Δ*hylB* (n = 18). CFUs were determined for liver, spleen, and kidney 24 HPI. Each symbol represents the CFU/gram of tissue recovered from a single mouse, and bars indicate the geometric mean. *P*_MW_ = Mann Whitney U Test of CFU data; *P*_Fe_ = Fisher’s exact test.

As we observed a reduced incidence of secondary bacteremia in the CAUTI model and *E. faecalis* is a common cause of CLABSI, we next sought to determine whether HylA and HylB contribute to primary bacteremia. CBA/J mice were inoculated via tail vein injection with 1 x 10^8^ CFU of V587, Δ*hylA*, Δ*hylB*, or Δ*hylA*Δ*hylB*, mice were sacrificed 24 HPI, and CFUs were quantified in the liver, spleen, and kidneys (Figure 4B). While the Δ*hylA* and Δ*hylB* mutants exhibited similar overall colonization to V587 in this model, the Δ*hylA*Δ*hylB* double mutant had a significant colonization defect in the kidneys and the spleen. Thus, HylA and HylB may be functionally redundant during primary bacteremia.

### Constitutive expression of *hylB* confers hyaluronidase and chondroitinase activity

With a contribution to pathogenesis confirmed, we next sought to define the enzymatic properties of HylA and HylB in *E. faecalis* V587. To bypass any issues due to endogenous gene expression, we opted to constitutively express *hylA* or *hylB* in the Δ*hylA*Δ*hylB* double mutant. HylB is predicted to be fully secreted and lacks complex domain architecture, so the full-length sequence of *hylB* was cloned under control of the *ermB* promoter in pAOJ84, and exported protein was detected as the expected molecular weight of 109 kDa (Figure 5A). HylA is predicted to be anchored to the cell wall, which presents a potential challenge for assays requiring unattached secreted enzyme and could also impact cell wall integrity when over-expressed. We therefore designed vector pAOJ86 to contain the full-length *hylA* sequence lacking the LPXTG anchor, and exported protein was again detected at the expected molecular weight of 146 kDa (Figure 5A).

**FIG 5.**
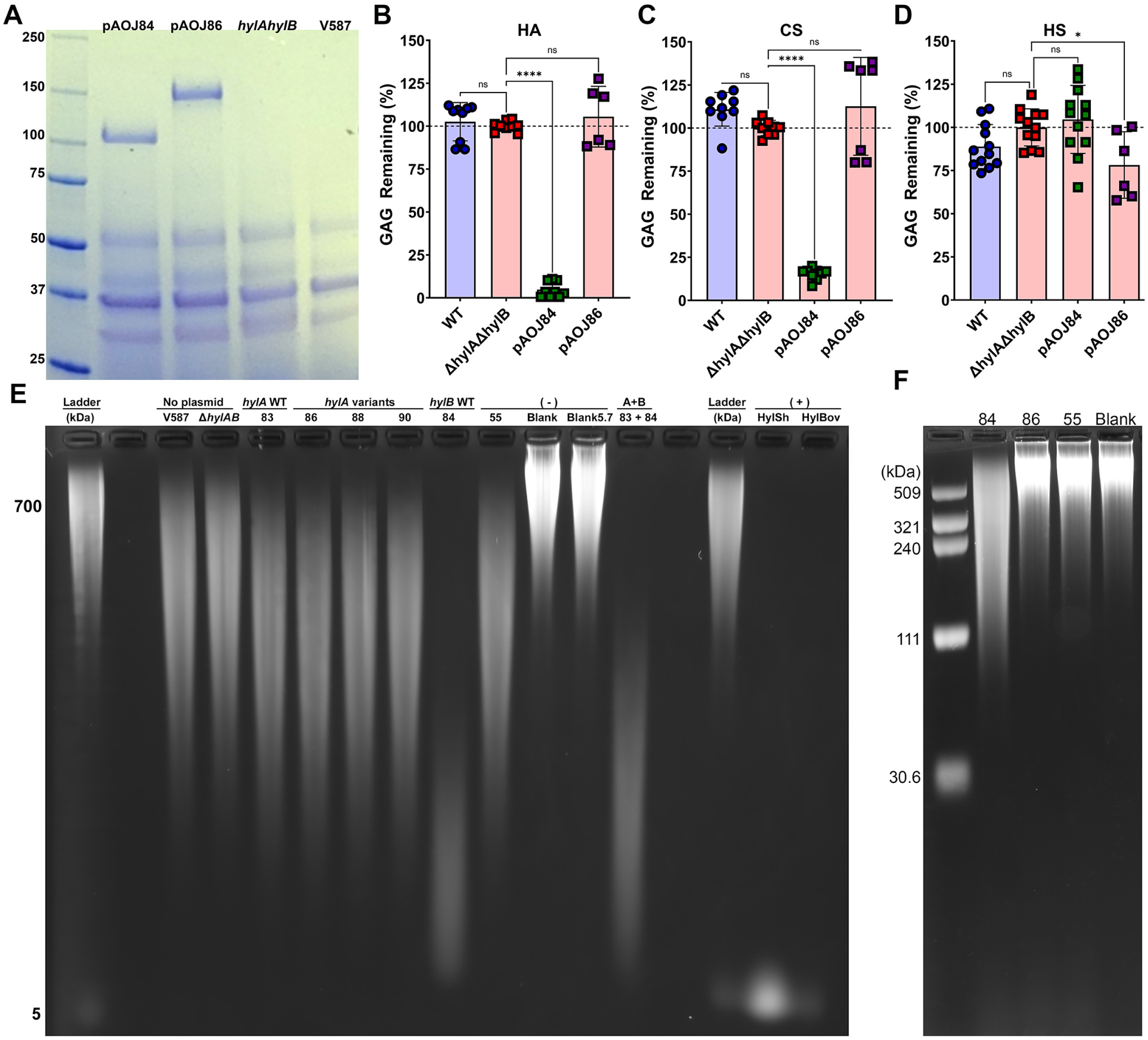
HylA and HylB expression and GAG degradation. (A) Coomassie Brilliant Blue stained SDS-PAGE gel of soluble secreted protein from cell-free supernatants of Δ*hylA*Δ*hylB* constitutive expressing *hylB* (pAOJ84), *hylA* (pAOJ86), or a control plasmid (pAOJ55), with V587 included as a control. Gel is representative of at least three independent experiments. (B-D) *E. faecalis* strains were incubated with 2.5 mg/mL of HA (B), CS (C), or heparin sodium salt (HS, D) in 0.5X TSB for 24 hours and supernatants were measured for remaining GAG by precipitation with BSA/acetic acid. Percentage GAG remaining was calculated by comparison to a curve of undigested GAG standard concentrations in bacteria-free samples, and data were normalized to Δ*hylA*Δ*hylB* for each individual experiment. Error bars indicate mean ± standard deviation (SD) for at least three independent experiments with two replicates each. \**P*<0.05,\*\*\*\**P*<0.0001 by one-way ANOVA with Dunnett’s multiple comparison correction. (E) *E. faecalis* strains were incubated in BHI with 1 mg/mL HA for 24 hours and HA degradation was examined by gel electrophoresis. The ladder was generated by combining 700 kDa and 5 kDa HA. Labels above each lane indicate *E. faecalis* strain, with the Δ*hylA*Δ*hylB* strains containing expression constructs abbreviated by plasmid number, in which 55 refers to the vector control. “Blank” indicates uninoculated BHI with HA and “5.7” indicates BHI with HA adjusted to pH 5.7. “83 + 84” indicates inoculation with an equal amount of Δ*hylA*Δ*hylB* (pAOJ83) and Δ*hylA*Δ*hylB* (pAOJ84). HylSh and HylBov are *Streptomyces* and bovine hyaluronidase, respectively. (E) The indicated *E. faecalis* strains were grown in BHI for 24 hours at 37°C, then filter-sterile supernatant was concentrated on a 10 kDa mol wt cutoff filter, and the retentates were incubated with 1 mg/mL HA in reaction buffer for 24 hours before subjecting the samples to electrophoresis.

Using an established semi-quantitative method for measuring GAG digestion with live bacteria (29), we incubated *E. faecalis* WT, Δ*hylA*Δ*hylB*, and Δ*hylA*Δ*hylB* containing the constitutive expression vectors for 24 hours at 37°C under low oxygen conditions in 0.5X TSB containing 2.5 mg/mL of hyaluronic acid (Figure 5B). In agreement with the literature and our prior results, V587 did not digest any of the GAGs and had an identical profile as the negative control strain Δ*hylA*Δ*hylB* (Figure 5B-D), confirming that *E. faecalis* does not exhibit GAGase activity under *in vitro* conditions (29). Unexpectedly, Δ*hylA*Δ*hylB* expressing *hylA* (pAOJ86) did not exhibit any HA degradation, but Δ*hylA*Δ*hylB* expressing *hylB* (pAOJ84) fully degraded HA, confirming that HylB is indeed a hyalouronidase.

Four additional variants of HylA were generated for constitutive expression to account for potential issues in protein folding, including 1) full-length HylA with the LPXTG anchor intact (pAOJ83), 2) HylA lacking both the LPXTG and FIVAR/repeat domains (pAOJ87), 3) HylA lacking the non-enzymatic conserved domains at the N-terminus (pAOJ88), and 4) a HylA chimera in which the LPXPTG anchor was removed and the N terminal region upstream of the predicted GAGase domain was replaced with the N terminal region upstream of the GAGase domain of HylB (pAOJ90). All resulted in secreted protein at the correct molecular weight except for pAOJ87 (Supplemental Figure 1), which may indicate a requirement for the FIVAR/repeat domain for proper protein expression, stability, or secretion. However, none of the HylA expression vectors conferred HA degradation, indicating that either none of the HylA constructs produce catalytically-active enzyme or that HylA acts on a different substrate.

To determine if either HylA or HylB act on other GAGs, we repeated the assay using chondroitin sulfate (CS, Figure 5C) and heparin sodium salt (HS, Figure 5D). The strain expressing HylB degraded the majority of the CS, which was not entirely unexpected as hyaluronidases from other bacteria have been shown to exhibit activity against CS (41). In contrast, the only phenotype observed for the strain expression HylA was a slight though statistically significant decrease in HS levels.

An important consideration when studying GAG degradation is the final product size. Host hyaluronidases and other factors such as oxidative damage (42) can cleave HA into intermediate sized fragments (< 500 kDa) or oligosaccharides that act as damage-associated molecular patterns and stimulate inflammation via TLR-2 and/or TLR-4 (10, 43, 44), while hyaluronidases from some pathogenic bacteria can cleave terminal disaccharides from HA which are anti-inflammatory (18, 45). Since the semi-quantitative GAG degradation assay is based on precipitation of high molecular weight GAGs, it would not detect subtle decreases in GAG fragment size. We therefore incubated HA directly with live *E. faecalis* V587, Δ*hylA*Δ*hylB* or Δ*hylA*Δ*hylB* harboring each of the constitutive hyaluronidase constructs or a vector control (pAOJ55) for 24 hours and used agarose gel electrophoresis to examine HA degradation product size (Figure 5E). An uninoculated control was incubated under the same conditions. Since *E. faecalis* acidifies BHI to a pH of ∼5.7 after 24 hours, an additional BHI control was adjusted to pH 5.7 with lactic acid to account for any HA degradation that might occur due to acidification alone. Incubation of HA with commercially available hyaluronidases from *Streptomyces hyalurolyticus* (HylSh) and Bovine Testicular Hyaluronidase Type I-S (BovHyl), that both digest HA down to short oligosaccharides (46), were utilized as positive controls.

Incubation with HylSh and BovHyl fully degraded the HA to a small band at ∼5 kDa, while no degradation was observed for either BHI control (Figure 5E). Importantly, HylSh is highly specific for HA (46), confirming the identity of the bands stained by this method. The strain containing pAOJ84 (*hylB*) displayed obvious digestion of HA and left only a low molecular weight smear of degradation products visible on the gel, albeit larger than the ∼5 kDa products of HylSh or BovHyl (Figure 5E). No degradation was observed for any of the HylA constructs. We hypothesized that HylA may only act on lower molecular weight HA fragments produced through the action of HylB, and further digest them to even smaller oligo- or disaccharides. To test this hypothesis, we incubated HA with a combination of Δ*hylA*Δ*hylB* carrying pAOJ83 (full length *hylA*, in case the LPXTG cell wall anchor was required for full activity) and Δ*hylA*Δ*hylB* carrying pAOJ84 (*hylB*). The combination still only resulted in a low molecular weight smear of HA similar to that of Δ*hylA*Δ*hylB* carrying pAOJ84 alone (Figure 5E), indicating that HylA does not provide any additional depolymerization of HA digested by HylB.

A puzzling observation was that all *E. faecalis* strains including Δ*hylA*Δ*hylB* partially depolymerized HA, resulting in a large smear that ran lower on the agarose gel than the BHI negative controls at either neutral pH or pH 5.7 (Figure 5E). This observation suggests that *E. faecalis* V587 has an additional mechanism to partially degrade HA that would not have been apparent in the semi-quantitative assay (Figure 5B) as large fragments would still precipitate and appear as undigested HA. To determine if *E. faecalis* exported protein was responsible for the observed basal level of HA degradation, HA was incubated with cell-free supernatants of *E. faecalis* Δ*hylA*Δ*hylB* carrying pAOJ84 (*hylB*), pAOJ86 (*hylA*), or pAOJ55 (vector control) that had been filter sterilized and concentrated using a 10 kDa molecular weight cutoff filter (Figure 5F). Concentrated supernatants from the strain constitutively expressing *hylB* showed partial digestion of high molecular weight HA, possibly suggesting that continual production of HylB is required for reaction completion. However, no HA degradation was observed for any of the other strains, indicating that the partial HA degradation that occurs during incubation with live bacteria is not mediated by a soluble, secreted product ≥10 kDa.

Intrigued by this result, we searched the genome of V587 and the related V583 for other potential GAGases and noted the presence of a putative heparinase, EF2268, that could be responsible for the weak digestion. No signal peptide was predicted in the EF2268 amino acid sequence by SignalP 5.0, which would align with our observation that HylB was the only large, secreted product that depolymerized HA (Figure 5E). We therefore deleted *ef2268* from the Δ*hylA*Δ*hylB* strain background to generate a triple GAGase mutant. Surprisingly, partial depolymerization was still observed during incubation with the triple mutant, demonstrating that EF2268 is not responsible for the basal HA degradation (Supplemental Figure 2).

### HylB purified from *E. coli* retains hyaluronidase activity

To characterize the activity of purified HylB enzyme, we cloned *hylB* into the *E. coli* expression plasmid pBAD/myc his A, transformed the plasmid into *E. coli* BL21(DE3) pLysS, and purified the recombinant HylB-Myc-6x His protein. The extracted protein was of high purity (Supplemental Figure 3). While *E. coli* BL21(DE3) pLysS is not suspected to encode any enzymes that could degrade HA, we used lysate from a culture of *E. coli* expressing unrelated proteins in a pBAD/myc his A background to serve as a negative control. Hyaluronidase activity was measured in a turbidimetric assay designed for use with purified protein rather than live bacteria. In this assay, HylB-Myc-6x His achieved complete HA digestion in less than 15 minutes, compared with ∼40% HA remaining with bovine hyaluronidase at equimolar protein concentration (Figure 6A). Digestion product size was also examined by agarose gel electrophoresis. Incubation of a 601 kDa HA fragment with *Streptomyces* hyaluronidase (HylSh) completely digested all HA, such that no remaining product was visible on the gel (Figure 6B). Incubation with recombinant HylB-Myc-6x His resulted in near complete HA digestion with only a low molecular weight smear around ∼30 kDa remaining, mirroring the results obtained with live *E. faecalis* cells (Figure 6B). Importantly, incubating HA with the *E. coli* protein lysate showed no digestion in either the gel or turbidometric assay (Figure 6A and B), confirming that all activity was derived from HylB-Myc-6x His. Combined, these data confirm HylB as a hyaluronidase, and demonstrate that purified HylB enzyme is sufficient for complete digestion of HA. While a similar approach was tried for expression and purification of HylA, all attempts were unsuccessful. Thus, the function of HylA remains to be defined.

**FIG 6.**
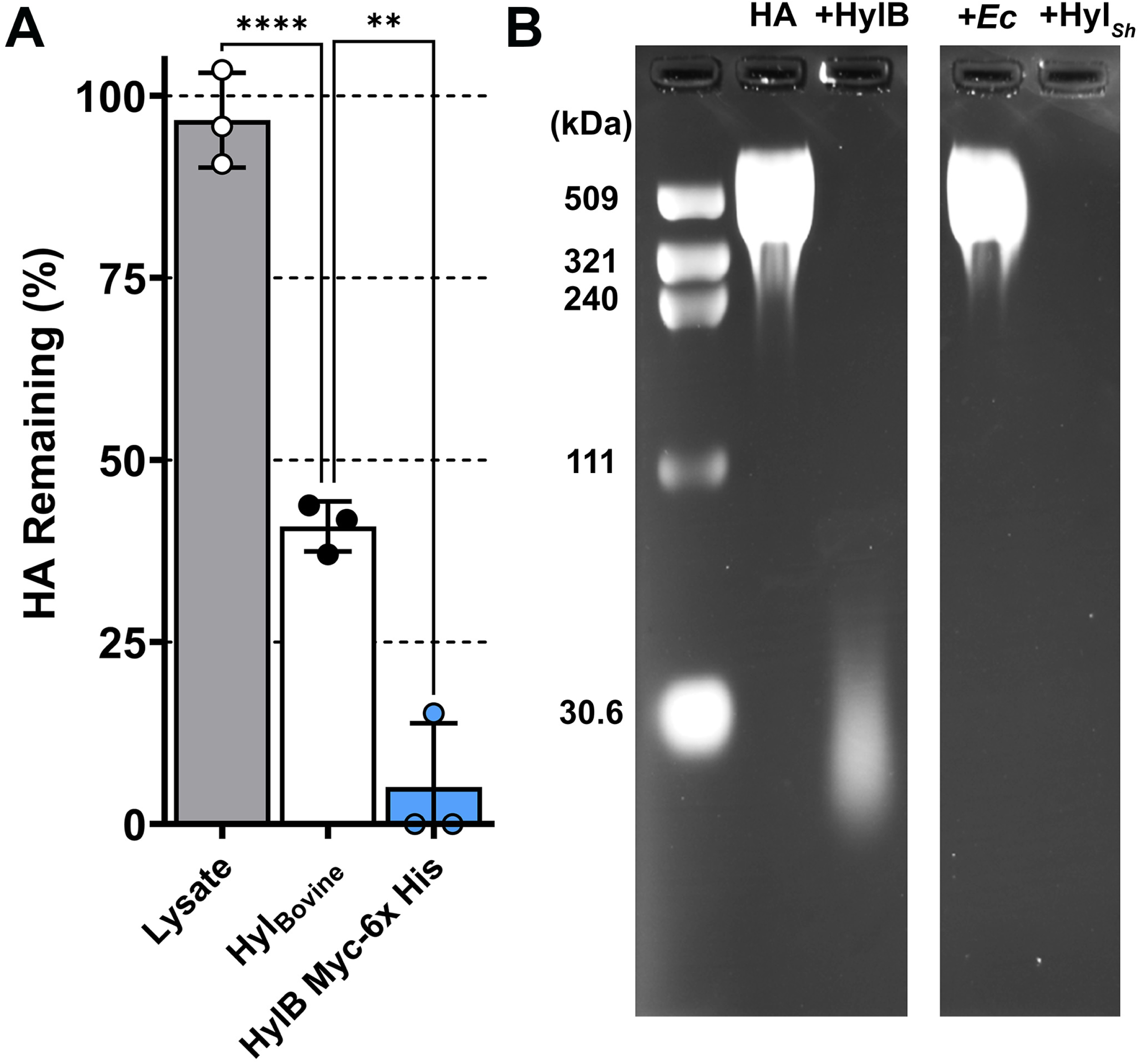
Characterization of purified HylB. (A) Turbidimetric assay of the remaining amount of HA post treatment with *E. coli* protein lysate, Bovine Hyaluronidase Type I-S, or HylB-Myc-6x His overexpressed and purified from *E. coli* after a 15 minute incubation at 37°C. Protein concentrations were adjusted based on molecular weight to obtain approximately equimolar amounts to 12.5 µg/mL Bovine Hyaluronidase, with the exception of the mixed soluble *E. coli* proteins which were used at a concentration of 50 µg/ml. The percentage of HA remaining is based on the BSA negative control. Error bars indicate mean ± SD for three experimental replicates. (B) Agarose gel electrophoresis of a 601 kDa HA fragment digested with recombinant HylB. The 601 kDa HA fragment was incubated overnight at 37°C with either: assay buffer and no enzyme (-); recombinant HylB-Myc-6x His (HylB); soluble *E. coli* protein; or commercially purchased *Streptomyces* hyaluronidase (HylSh). Image was captured on a ChemiDoc MP, irrelevant lanes were cropped out as indicated by tool marks, and the levels were adjusted for better visibility with Adobe Photoshop 2022.

### *E. faecalis* does not use HA as a nutrient source during growth *in vitro*

Some bacteria can use HA as a nutrient source after breaking down the repeating polymers into N-acetyl-glucosamine and glucuronic acid monomers. For example, the hyaluronate lyase activity of *S. pneumoniae* allows this species to use HA as a carbon source in minimal media (21). We therefore sought to determine whether HA degradation by HylB could support growth of *E. faecalis* in a simplified defined media (SDM) (47) in which *E. faecalis* is unable to grow without a supplied carbohydrate source (47). *E. faecalis* V587 exhibited robust growth in SDM supplemented with glucose, but failed to grow in SDM with HA as the sole carbohydrate (Figure 7A). However, HylB may not be active in WT *E. faecalis* under these conditions, even when HA is supplied as the sole carbohydrate source. Since HylB-mediated degradation of HA was clearly observed in the semi-quantitative GAG degradation assay in 0.5X TSB, we next asked whether HA supplementation enhanced growth when HylB was constitutively expressed. V587, Δ*hylA*Δ*hylB*, and Δ*hylA*Δ*hylB* with pAOJ84 (constitutively expressing *hylB*) all grew similarly in 0.5X TSB (Figure 7B) and they all achieved stationary phase at a similar rate and density when supplemented with HA (Figure 7C). The addition of unfractionated HA caused a slight reduction in the cell density at which stationary phase was achieved for all strains. However, constitutive expression of *hylB* resulted in an increase in Δ*hylA*Δ*hylB* culture density from 8-24 hours that was not observed for the other strains (Figure 7C), indicating that HylB activity either provides *E. faecalis* with an additional nutrient source or alleviates any inhibition imposed by the presence of high molecular weight HA.

**FIG 7.**
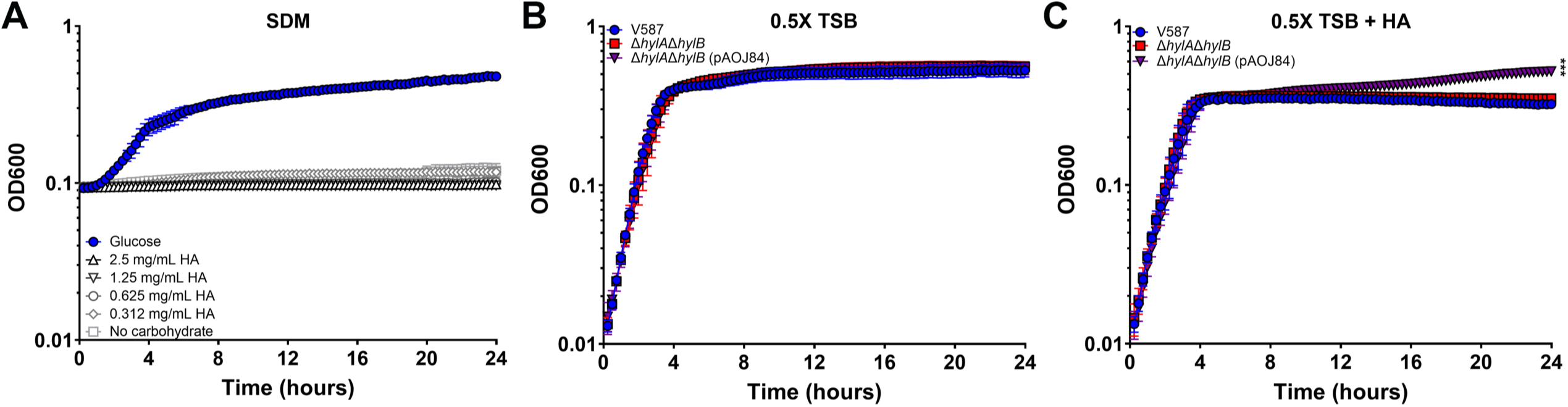
Contribution of HA degradation to *E. faecalis* growth. (A) *E. faecalis* V587 was incubated in simplified defined medium (SDM) supplemented with either12 mM glucose or a series of HA concentrations and growth was measured by OD_600_ every 15 minutes for 24 hours. Graph is representative of two independent experiments. Error bars indicate mean ± SD for 10 technical replicates. (B-C) *E. faecalis* V587, Δ*hylA*Δ*hylB*, and Δ*hylA*Δ*hylB*, expressing *hylB* (pAOJ84) were incubated in 0.5X TSB (B) or 0.5X TSB supplemented with 2.5 mg/ml of unfractionated high molecular weight HA (C). Growth was measured by OD_600_ every 15 minutes for 24 hours. Graph is representative of two independent experiments. Error bars indicate mean ± SD for 10 technical replicates. \*\*\**P*<0.001 by two-way ANOVA with Dunnett’s multiple comparison correction.

### HylA and HylB contribute to suppression of LPS-induced inflammation but do not dampen inflammation during CAUTI

Hyaluronidases from other Gram-positive bacteria have been shown to mediate inflammatory regulation (18). For example, *S. agalactiae* digests HA to disaccharides that facilitates pathogenesis by interfering with TLR-2/TLR-4 signaling (18) and subverting neutrophil and macrophage killing (19, 48, 49). Since *E. faecalis* is well-known for the ability to modulate the immune response, including suppression of lipopolysaccharide (LPS) stimulated inflammation (50, 51), we sought to determine whether HylA and HylB contribute to modulation of the innate inflammatory response using a well-established NF-κB macrophage-like reporter cell line, RAW-Blue. This cell line expresses a secreted embryonic alkaline phosphatase (SEAP) under the control of an NF-κB inducible promoter, allowing for a readout of NF-κB induction via an alkaline phosphatase assay.

To determine if HA disaccharides dampen NF-κB activation in this cell line, RAW-Blue cells were incubated for six hours with increasing concentrations of HA disaccharides, LPS, and each combination thereof (Figure 8A). Interesting, the HA disaccharides appears to be pro-inflammatory at 0 and 0.1 ng/ml of LPS, but dampened NF-κB induction at higher concentrations of LPS. While these effects were all modest, the data confirm that HA disaccharides can dampen LPS-stimulated activation of NF-κB in RAW-Blue cells.

**FIG 8.**
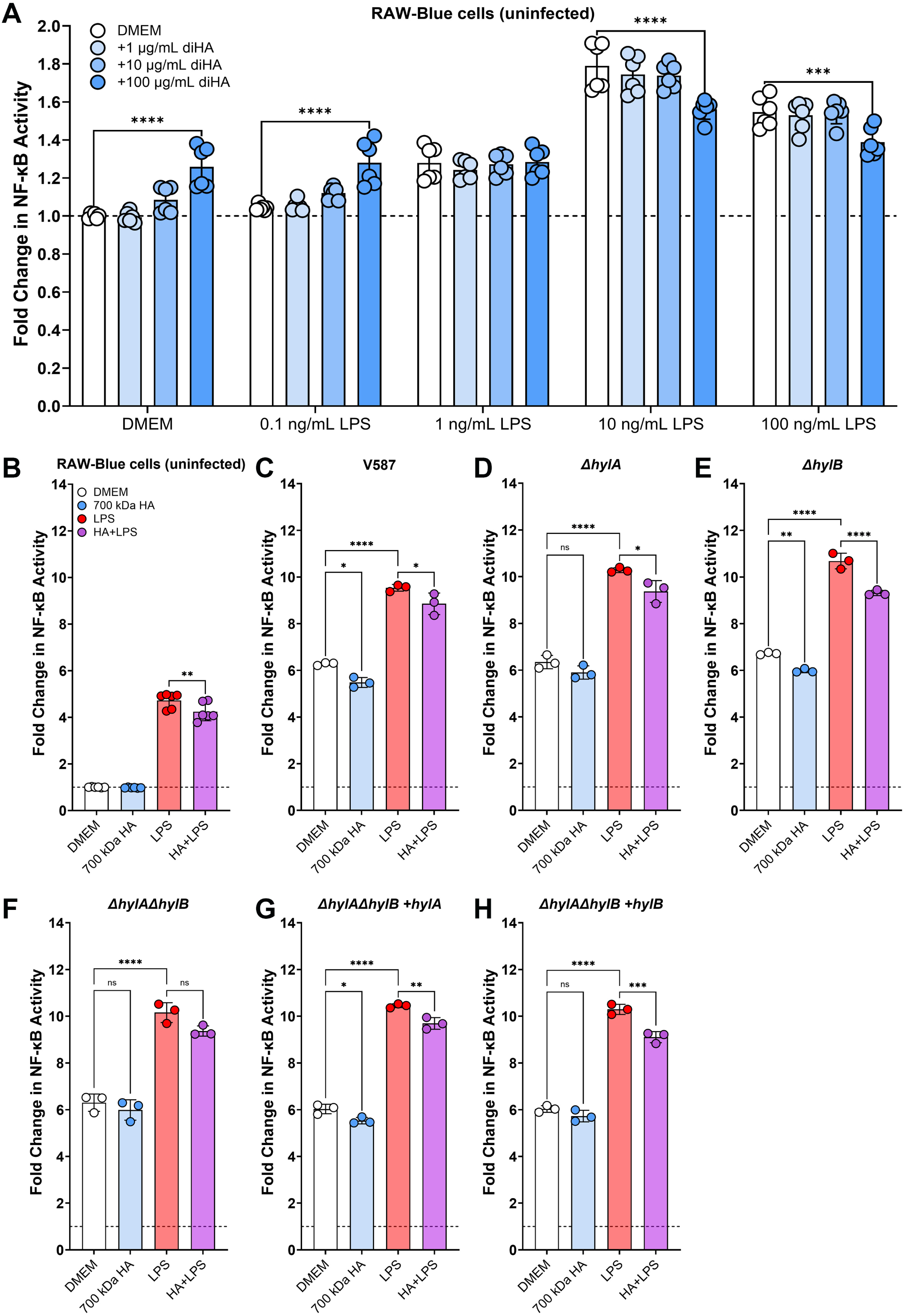
HA degradation reduces LPS-stimulated NF-κB activation. (A) RAW-Blue cells were grown in a 96-well plate in Dulbecco’s Modified Eagle Medium (DMEM) and stimulated with increasing concentrations of LPS with or without HA disaccharides. NF-κB activity was measured by transferring 20 µL of supernatant to 180 µL alkaline phosphatase detection medium, incubating with shaking at 37°, then reading the OD_640_ nm at 2 hours post incubation. Error bars display mean ± SD from two independent experiments with three replicates each. \*\*\*\**P*<0.0001 by two-way ANOVA with Dunnett’s multiple comparison correction. (B-H) RAW-Blue cells were incubated alone or infected for six hours with the indicated *E. faecalis* strain at an MOI of 10 in DMEM with or without 100 ug/ml of 700 kDa HA and 10 ng/ml LPS. Error bars display mean ± SD from one independent experiments with three replicates and are representative of two independent experiments. \**P*<0.05, \*\**P*<0.01, \*\*\**P*<0.001,\*\*\*\**P*<0.0001 by one-way ANOVA with Dunnett’s multiple comparison correction.

We next examined NF-κB activation in response to infection. In preliminary experiments, we observed that infection with MOIs greater than 10 resulted in substantial acidification of the culture media before assay completion. Since low pH has been shown to interfere with the NF-κB pathway (52–54), an MOI of 10 was chosen for all experiments. RAW-Blue cells were incubated for six hours with wild-type *E. faecalis* V587, Δ*hylA*, Δ*hylB*, Δ*hylA*Δ*hylB*, Δ*hylA*Δ*hylB* with pAOJ83 (constitutively expressing the full length HylA protein), or Δ*hylA*Δ*hylB* with pAOJ84 (constitutively expressing the full length HylB protein) with or without 10 ng/mL LPS and 100 ug/ml of high molecular weight HA (∼700 kDa). In uninfected cells, LPS supplementation resulted in a ∼5-fold increase in NF-κB activation as expected while the addition of HA had no impact under any conditions (Figure 8B), confirming that 700 kDa HA does not alter stimulation in the absence of hyaluronidase activity.

Infection with any of the *E. faecalis* strains resulted in a >6-fold increase in NF-κB activation over DMEM alone (Figure 8C-H), clearly demonstrating that strain V587 is capable of inducing inflammation at an MOI of 10. The addition of HA during infection with V587 resulted in a slight though statistically-significant decrease in NF-κB activation, which was also observed during infection with Δ*hylB* or Δ*hylA*Δ*hylB* constitutively expressing the full length HylA protein but not during infection with any strain lacking *hylA*. These data suggest that HylA may provide a small degree of immune suppression in the absence of other stimuli.

Supplementation with LPS during infection resulted in much greater NF-κB activity for all strains compared to infection in DMEM alone (∼10-fold induction vs ∼6-fold), demonstrating that *E. faecalis* V587 does not suppress LPS-stimulated NF-κB activation under these experimental conditions. This is in contrast to prior reports and likely due to differences in MOI and subsequent media acidification. The addition of HA slightly mitigated NF-κB activation during infection, as HA+LPS resulted in a modest though significant decrease in NF-κB activation for all strains except the Δ*hylA*Δ*hylB* double mutant. Since loss of either gene alone or complementation with either gene alone allowed for decreased in NF-κB activation, these data suggest that digestion of HA by either HylA or HylB is sufficient to modestly suppress LPS-stimulated inflammation.

We next sought to determine whether HylA or HylB contribute to immune modulation during CAUTI. Since individual mutants exhibited colonization defects 48 HPI in the CAUTI model, we conducted three additional infection studies with V587 and the Δ*hylA*Δ*hylB* mutant to examine colonization at 6 HPI and two studies at 24 HPI (Figure 9A and B). No significant differences in colonization were observed at these earlier time points, allowing us to pinpoint the potential contribution of HylA and HylB to immune modulation.

**FIG 9.**
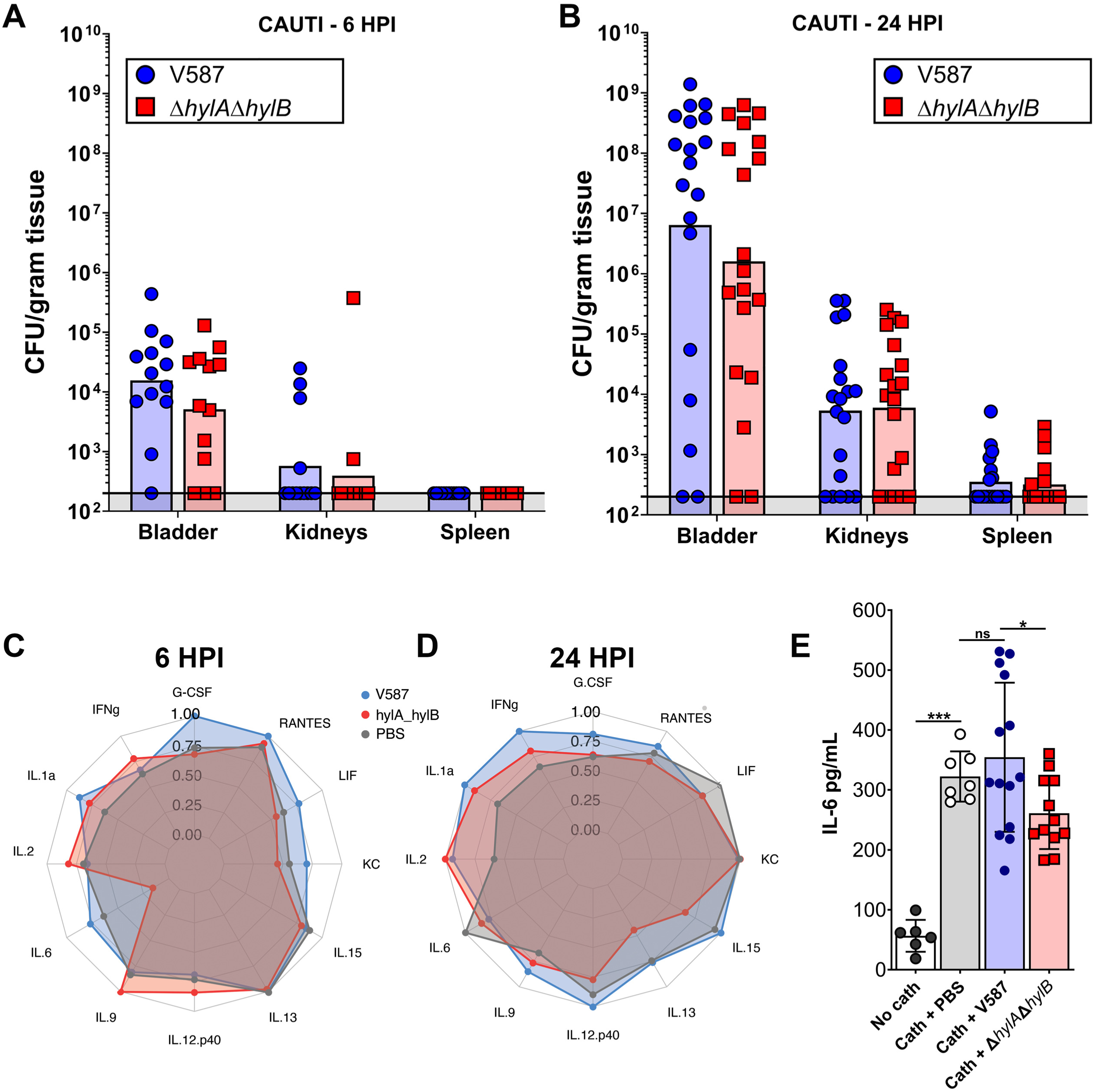
Contribution of HylA and HylB to bladder cytokine and chemokine profiles. (A-B) 7-week old female CBA/J mice were inoculated via transurethral catheterization with 2 x 10^5^ CFU of V587 or Δ*hylA*Δ*hylB*, or inoculated with a PBS control then sacrificed at 6 and 24 HPI. Each data point represents the CFU/g obtained from a single mouse, and bars indicate the geometric mean. (C-D) Radar graphs for select cytokines and chemokines in bladders from mice inoculated with V587, Δ*hylA*Δ*hylB* , or PBS at (C) 6 hours and (D) 24 hours post-inoculation. Each ring in the graph represents quartiles of the given analyte, with the furthest ring representing the upper bound of the range of the log of mean abundance (pg/ml) for each metabolite. The scales between A and B are the same for ease of interpretation and the max values (outer-ring) for each metabolite is provided in the legend. (E) IL-6 levels were quantified by ELISA in bladder homogenates from mice that were either mock-inoculated with PBS, catheterized and mock-inoculated with PBS, or catheterized and infected with either V587 or Δ*hylA*Δ*hylB*. Error bars represent mean ± SD. \**P*<0.05, \*\*\**P*<0.001 by one-way ANOVA with Dunnett’s multiple comparison correction.

For initial assessment of the innate immune response, bladder homogenates were pooled by experiment date and infection group, resulting in three biological samples per inoculum at 6 HPI and two biological samples at 24 HPI. Homogenates from mock-infected (PBS) catheterized mice were also included for each time point. Samples were then analyzed by Luminex to quantify the levels of 32 cytokines, chemokines, and growth factors, of which 14 were above the limit of detection (Figure 9C and D, Supplemental Figure 4). Unexpectedly, the global profiles of the catheterized mock-infected PBS mice were not significantly different from either infection group at either time point, suggesting that the majority of the inflammation was driven by the presence of the catheter. This, *E. faecalis* V587 does not appear to substantially dampen catheter-driven inflammation, and also does not further stimulate the immune response in this model.

To determine the potential contribution of HylA and HylB to inflammation, the profile of bladders from mice infected with V587 were compared to those infected with. Interestingly, the only analyte that was significantly different at any time point was a reduction in IL-6 at 6 HPI for mice infected with the Δ*hylA*Δ*hylB* double mutant (*P* = 0.0004). This observation was unexpected, as we hypothesized that HylA and HylB would most likely dampen the immune response, resulting in an increase in pro-inflammatory cytokines during infection with the mutant strain. To confirm whether HylA and HylB might contribute to IL-6 induction, an additional set of mice were inoculated as above along with an extra control group lacking the catheter segment, and IL-6 was quantified by ELISA (Figure 9E). As expected, the presence of the sterile catheter segment alone resulted in ∼8-fold induction of IL-6 in the absence of infection, but no further increase in IL-6 was observed during infection with V587. These data confirm the hypothesis that the catheter segment drives the majority of inflammation in this model at 6 HPI. Further, IL-6 levels were significantly lower in mice inoculated with the Δ*hylA*Δ*hylB* double mutant compared to V587, confirming that HylA or HylB contribute to stimulation of IL-6. It is important to note that the effect is modest at 6 HPI and appears to be resolved by 24 HPI, and likely does not substantially contribute to the differences in dissemination that are observed in this model. In summary, HylA and HylB may contribute to immune modulation but do not appear to be the primary mediators of immune suppression during CAUTI.

## Discussion

*Enterococcus faecalis* infections are very common and impose a high burden on the healthcare system, ranking as the 5^th^ most commonly encountered healthcare acquired pathogen in the United States overall, both in pediatric patients and adults (1, 2). *E. faecalis* can cause a variety of infections, including CAUTI and CLABSI (2, 3). In this study, we revealed HylA and HylB as two new virulence factors for CAUTI and bacteremia caused by vancomycin-resistant *E. faecalis*.

Even though we detected *hylA* and *hylB* mRNA transcripts in WT *E. faecalis* V587 during growth *in vitro,* we did not detect any hyaluronidase activity in the WT strain. These data confirm prior work indicating a lack of HA degradation by *E. faecalis.* However, by cloning *hylB* under a constitutive promoter in *E. faecalis* strain V857 we conclusively demonstrated that HylB (EF0818) is a bona fide hyaluronidase with substantial chondroitinase activity. This enzyme appears to act specifically on HA and CS, as we did not detect any evidence of HS digestion. We further demonstrated that expression and purification of HylB in *E. coli* also provides robust hyaluronidase activity, showing that HylB is both necessary and sufficient to digest HA. As the amino acid sequencing of HylB in *E. faecalis* V587 shows low identity with previously characterized Gram-positive bacterial hyaluronidases (55–59), structural studies of this enzyme could provide critical information for GAGases in general.

In contrast to HylB, the only *in vitro* evidence of enzymatic activity for HylA (EF3023) was a very subtle degradation of HS and a potential role in dampening NF-κB activation in the presence of HA. However, *hylA* contributed to bladder colonization and dissemination to the bloodstream in the mouse model of CAUTI and appeared to provide functional redundancy to *hylB* in the bacteremia model. There are many potential explanations for the discrepancy between the *in vitro* and *in vivo* phenotypes of HylA. One possibility is that none of the HylA expression constructs resulted in properly folded or active enzyme. Alternatively, HylA may require processing by additional factors not present under our *in vitro* conditions to become a catalytically-active enzyme. For example, the HylB enzyme of *S. agalactiae* requires post-translational proteolytic cleavage to be active (60). Another possibility is that the substrate of HylA may be a specific glycosaminoglycan, glycoprotein, or sulfonation pattern that was not directly tested in this study but is present in the urinary tract and circulatory system. This would not be the first time that a predicted bacterial hyaluronidase turned out to have a different target substrate; *Streptococcus pyogenes* Spy1600 was initially labeled as a possible hyaluronidase but experimentally confirmed to act as a β-N-acetylglucosaminidase (61). Experimentation with additional host polysaccharides and expression constructs will be needed to either confirm or disprove HylA as a glycosaminoglycan degrading enzyme.

While some bacteria such as *S. pneumoniae* can use HA as a carbon source (21), this was not the case for *E. faecalis in vitro*. However, constitutive expression of *hylB* did allow the Δ*hylA*Δ*hylB* double mutant to achieve higher cell density during HA supplementation than even the WT strain. Considering the proximity of *hylB* to a putative oligosaccharide PTS system, there is likely additional regulation that must occur for *E. faecalis* to potentially use HA degradation products as a nutrient. Our data also do not rule out a possible role for HA catabolism to fuel growth during infection.

The contribution of hyaluronidases to microbial pathogenesis has been studied in *S. agalactiae* in the context of pre-term birth (48, 49) and systemic infection (19). During systemic infection in mice, disruption of hyaluronidase by mutation of *hylB* decreased tissue colonization and lethality (19). In a nonhuman primate model, hyaluronidase activity decreases neutrophil bactericidal activity by interfering with TLR2/TLR4 signaling, and a *hylB* mutant deficient in hyaluronidase activity was deficient in establishing fetal invasion and bacteremia (48). In mice, the *hylB* mutant colonized the vaginal tract to a similar level as the parental isolate but exhibited defects in ascending infection and in the immune-subversion (19). Vaginal colonization has also been examined in *E. faecalis* strain OG1RF using a genome-wide transposon mutant screen for factors that allow *E. faecalis* to persist within the vaginal tract (62). Intriguingly, the homolog of *hylB* in this strain (OG1RF_10550) exhibited reduced colonization at all time points (62).

However, OG1RF_10550 was just one of many genes identified as important for vaginal colonization in this study, and we are unaware of any follow-up studies on this gene. Considering that our studies demonstrate that *E. faecalis* HylA primarily contribute to bladder colonization and dissemination to the bloodstream while HylB contributes to kidney colonization and almost 50% of *E. faecalis* isolates possess homologs of both genes, expression and activity are likely highly regulated. This is supported by the recent observation that HylA expression is regulated by a BglG/SacY antiterminator homolog in *E. faecalis* strain V19, a plasmid-cured derivative of V583 (63). Further exploration into the conditions and signaling pathways required for each to be expressed and active is expected to provide further insight into how *E. faecalis* is able to adapt to many different infection niches.

Hyaluronic acid also modulates inflammatory signaling, with the outcome being largely dependent on HA fragment size. Hyaluronidases from *S. agalactiae* and other Gram-positive pathogens were shown to digest HA down to disaccharides, which block TLR-2 and TLR-4 signaling (18). In contrast, *Streptomyces* hyaluronidase produces HA oligosaccharides that stimulate inflammation, as do mammalian hyaluronidases (18). Our data indicate that infection of RAW-Blue cells with *E. faecalis* V587 activates NF-κB regardless of condition or infecting strain, but constitutive expression of HylB can modestly counteract LPS-stimulated activation of NF-κB. Thus, HA degradation by *E. faecalis* V587 could contribute to immune modulation during infection.

We also examined a broad array of cytokines, chemokines and growth factors in the bladders and the kidneys of mice infected via our CAUTI model. Strikingly, we did not observe drastic differences between infected and mock infected catheterized mice, which led us to confirm that the majority of inflammation during CAUTI with *E. faecalis* V587is due to the insertion of the catheter itself. We also did not observe dramatic difference in immune response between infection with wild-type V587 and the Δ*hylA*Δ*hylB* strain, other than a slight reduction in IL-6 in the bladders of mice infected with Δ*hylA*Δ*hylB* at 6 HPI. Infection with *E. faecalis* as well as catheter implantation are both known to stimulate IL-6 (64–66), so this observation may suggest a role for HylA or HylB in tissue invasion and damage.

In summary, this study highlights the importance of *in vivo* validation of *in silico* functional predictions. We have confirmed HylB as a hyaluronidase/chondroitinase enzyme in vancomycin-resistant *E. faecalis*, despite having low amino acid identity to other bacterial hyaluronidases. We have further demonstrated that active HylB can be produced *in vitro*, both by *E. faecalis* and by *E. coli,* and that *hylA* and *hylB* both contribute to *E. faecalis* pathogenesis. Further examination of the substrate and functions of HylA and the distribution of its target in mammalian hosts may reveal novel host-pathogen interactions or new information about GAG distribution in mammalian tissues.

## Materials and Methods

### Animal model ethics statement

Animal protocols were approved by the Institutional Animal Care and Use Committee of the University at Buffalo (MIC31107Y), in accordance with the Office of Laboratory Animal Welfare, the U.S. Department of Agriculture, and the Association for Assessment and Accreditation of Laboratory Animal Care. Mice were anesthetized with a weight-appropriate dose (0.1 ml for a mouse weighing 20 g) of ketamine/xylazine (80 to 120 mg/kg ketamine and 5 to 10 mg/kg xylazine) by intraperitoneal injection and euthanized by CO_2_ with vital organ removal.

### Bacterial strains and culture conditions

Vancomycin resistant *Enterococcus faecalis* strain V587 (NR-31979) (31) was obtained through BEI Resources, NIAID, NIH. This strain was chosen for its clinical history that mirrors our mouse infection models, and for containing a large number of known *E. faecalis* virulence genes (67), including Enterococcal Surface Protein (*esp*) previously implicated in urinary tract colonization (68). *E. faecalis* V587 and isogenic mutants were routinely cultured from frozen glycerol stocks in 5 ml of Brain Heart Infusion (BHI) broth (Dot Scientific) or Tryptic Soy Broth (TSB) with 500 µg/mL gentamicin sulfate (ACROS Organics, catalog # 455310250) in tightly capped 14-ml polypropylene culture tubes at 37°C with shaking at 300 RPM. *E. faecalis* harboring plasmids were cultured with 20 µg/mL chloramphenicol (Research Products International, catalog # C61000) instead of gentamicin.

### RNA Extraction and qRT-PCR

*E. faecalis* V587 was grown overnight in BHI plus gentamicin 500 µg/mL, 0.6 mL aliquots of the overnight culture were centrifuged and washed once with PBS (pH 7.4), the cell pellet was resuspended in 0.6 mL of PBS, then diluted 1:100 in one of the following media: BHI, 0.5X TSB (TSB diluted to 50% concentration with sterile water), 0.5X TSB with 2.5 mg/mL ∼5 kDa size fractionated HA (Lifecore Biomedical, catalog # HA5K), 0.5X TSB with 2.5 mg/mL unfractionated HA sodium salt from *Streptococcus equi* (Sigma-Aldrich), filter sterilized human urine (Lee BioSolutions catalog # 991-03-P-FTD, lot # 01J5567), human urine with 2.5 mg/mL ∼5 kDa size fractionated HA, or human urine with 2.5 mg/mL unfractionated HA. 3 mL cultures were used for samples with rich media as the base, and 6 mL cultures were used for the samples grown in urine to account for lower growth density. Cultures were incubated with shaking for 4 hours at 37°C, pelleted by centrifugation, washed once with 1 mL of TE buffer (pH 8), then suspended in 50 µl RLT Buffer with BME (Qiagen RNeasy Mini Kit). 1.5 mL safe-lock microfuge tubes (Eppendorf) were filled halfway with 0.5mm glass disruption beads (RPI research Products International), the suspended pellet was added to the bead tube, and samples were homogenized using a Bullet Blender Gold (Next Advance) at max speed for 5 minutes.

After bead beating, 250 µl of RLT Buffer with BME was added and samples were vortexed to mix. A small hole was punched in the bottom of the tube with an 18G needle (BD Precision Glide), and the tube was immediately placed inside a 1.5 mL microfuge tube to collect the homogenate while leaving the beads behind. The collected homogenate was centrifuged at 21,000 rcf for 10 minutes at 4°C, supernatant was removed, and 1 volume of 70% ethanol was added and vortexed to mix. The sample was then transferred to an RNeasy Mini spin column for on-column DNase digestion kit (Qiagen), and RNA was eluted with 40 µl of RNase free water. A second DNase digestion was then performed off-column by adding 0.1 volume of 10X DNase buffer and 1 µl of DNase (Invitrogen) and incubating at 37°C for 30 minutes. The digestion was terminated by adding 0.1 volumes of DNase inactivation reagent, incubating at room temperature for 2 minutes, and centrifuging for 90 seconds at 8000 rcf. cDNA was synthesized using the iScript cDNA Synthesis Kit (BioRad) and qRT-PCR was performed using PCRBio SyGreen Blue Mix Lo-Rox (PCR Biosystems) and a BioRad CFX-Connect Real Time system. Data were normalized to *recA* as the reference gene and analyzed via the ΔΔCT method.

### Generation of *E. faecalis* deletion mutants

Markerless deletion mutants of *E. faecalis* V587 were constructed by allelic exchange using the plasmids designated in the Supplemental primer table. Roughly 1,000 base pairs upstream and downstream of the desired deletion location were amplified via PCR and assembled into allelic exchange vector pIMAY (Addgene plasmid # 68939, gifted by Tim Foster) (69) via either restriction digest and ligation or NEBuilder HiFi DNA Assembly Mastermix (NEB), as indicated in the Supplemental vector table. Assembled vectors were propagated in *E. coli* Top10 and verified a combination of restriction digest and sequencing (Plasmidsaurus).

Plasmids were electroporated into *E. faecalis*, recovered and plated at 30°C, and isolated colonies were struck onto pre-warmed BHI-chloramphenicol plates and incubated overnight at 37°C to induced plasmid integration (70). Chloramphenicol-resistant colonies were then struck on BHI agar containing 1 µg/mL anhydrotetracycline hydrochloride (Cayman Chemical catalog # 10009542) to induce expression of the counter selection marker, and incubated overnight at 30°C. Large, isolated colonies from these plates were then re-struck on anhydrotetracycline BHI plate and incubated at 37°C overnight to ensure loss of the plasmid backbone. Mutants were PCR verified by checking for the presence of the desired mutation, the absence of the original gene, the absence of the pIMAY backbone, and retention of natural V587 plasmids using the primers described in the Supplemental primer table. The *esp* gene was also amplified using esp11 and esp12 from a previous publication (71). Verified colonies were cultured in BHI-gentamicin broth overnight to generate glycerol stocks and PCR-verified a second time. The Δ*hylA*Δ*hylB* double mutant was generated by sequential deletion of *hylB* from the Δ*hylA* strain using the method described above, and was verified by PCR as described above and by sequencing of ∼1200 base pairs up and downstream of each gene (Eton Biosciences).

### Hyaluronidase and Chondroitinase plate assay

HA plates were made using a published recipe (18). A hyaluronic acid solution was first generated by suspending 300 mg unfractionated HA sodium salt and 8 g of bovine serum albumin fraction V at pH 5.2 (Sigma-Aldrich) in 200 mL of water and slowly adjusting to pH 7.5 using 0.1 M sodium hydroxide. The HA solution was then filter sterilized and warmed to 37°C. The agar base (10 g Noble agar, 10 g yeast extract, 30 g Todd-Hewitt broth powder, and 800 mL of RO water) was autoclaved, cooled to ∼56°C, supplemented with the warmed hyaluronic acid solution, and 16 mL was distributed into petri dishes. Once cooled, plates were inoculated with 10 µL of overnights cultures of *E. faecalis* or *L. lactis* and incubated overnight at 37°C. After overnight incubation, plates were cooled to ambient temperature and flooded with 2 M acetic acid to precipitate high molecular weight GAGs, and incubated for at least 10 minutes. The acetic acid was then aspirated and the plates were imaged on an Axygen Gel Documentation System (Corning) with blue light illumination.

### Mouse infection models

For establishment of CAUTI, 6-8 week old female CBA/J mice (The Jackson Laboratory, strain # 000656) were inoculated transurethrally with 50 µl of a suspension of 2 x 10^6^ colony forming units (CFU)/mL in PBS of either wild-type V587, Δ*hylA*Δ*hylB*, Δ*hylA,* Δ*hylB*, or mock infected using PBS. A 4 mm segment of silicone tubing was placed in the bladder during inoculation to simulate a urinary catheter, as previously described (40, 72). At 6, 24 or 48 hours post infection, mice were sacrificed, and the bladders, kidneys and spleens were homogenized in 5 mL tubes (Eppendorf) containing 500uL of 3.2mm stainless steel beads (Next Advance) and 1 mL of sterile PBS in a Next Advance 5 E Gold Bullet Blender. Organ homogenates were plated for CFU counts and an aliquot of each homogenate was flash frozen and stored at -80°C for cytokine and chemokine measurements as described below.

For establishment of bacteremia, 7 week old female CBA/J mice were inoculated via tail vein injection of 100 µl of a suspension of 5 x 10^8^ CFU/mL of either wild-type V587, Δ*hylA*, Δ*hylB* or Δ*hylA*Δ*hylB*. 24 hours post infection, the mice were sacrificed and livers, spleens and kidneys were harvested, homogenized and plated for CFU counts.

### Construction of constitutive expression vectors for *E. faecalis*

pAOJ20, an *E. coli*-Gram-positive shuttle vector described previously (73), was utilized as the backbone for plasmids pAOJ83 (full length HylA), pAOJ84 (full length HylB), pAOJ86 (HylA_[codons 1-1334]_, minus LPXTG domain), and pAOJ55 (vector control with a non-enzymatic 5’ fragment of unrelated protein AtlA (74, 75)). All plasmids have *hylA* or *hylB* expression driven by the constitutive *ermB* promoter (76). To construct plasmid pAOJ88 (HylA_[codons Δ29-246, Δ1335-1372]_, HylA minus the Discoidin/Big2 domains and the LPXTG domain), pAOJ86 was amplified via inverse PCR using phosphorylated primers AOJ_726 and AOJ_727, and circularized by ligation. Plasmid pAOJ90 (fusion construct with the HylB N-terminus [codons 1-215] and HylA C-terminus [codons 247-1334, Δ1335-1372]) was constructed by performing inverse PCR on pAOJ84 using primers AOJ_729 and AOJ_730 to obtain the plasmid backbone and first 216 codons of *hylB*, which was fused to codons 247-1334 of *hylA* amplified via AOJ_731 and AOJ_732 by digesting both DNA fragments with BspQI (NEB) and ligation using T4 DNA ligase.

Due to technical issues encountered while propagating the constitutive expression constructs in *E. coli,* all ligation products were transformed into *Lactococcus lactis* NZ9000 (MoBiTec catalog # VS-ELS09000-01) for plasmid amplification. *L. lactis* competent cells were made and transformed using the previously described Lithium acetate-Dithiothreitol (DTT) method (77). All constructs were sequence-verified via Plasmidsaurus before transformation into *E. faecalis* V587 Δ*hylA*Δ*hylB*.

### Secreted protein profiles of *E. faecalis* V587, Δ*hylA*Δ*hylB*, and constitutive expression strains

*E. faecalis* overnight cultures were diluted 1:200 into 50 mL conical tubes containing 20 mL BHI. Cultures were further supplemented with 20 µg/mL chloramphenicol for Δ*hylA*Δ*hylB* with pAOJ83, pAOJ84, pAOJ86, pAOJ87, pAOJ88, and pAOJ90, or 500 µg/mL gentamicin sulfate for V587 WT and V587 Δ*hylA*Δ*hylB*. The tubes were tightly capped and incubated with shaking for 4 hours, at which point the cells were pelleted and the supernatant filter sterilized using a 0.22 µm PES filter. The 4 hour time point was chosen to balance bacterial cell density while limiting contact time with the broad-spectrum *E. faecalis* proteases GelE and SprE, which exhibit peak expression in late exponential phase (78). 20 ml of filter-sterilized supernatant was placed on ice to cool, mixed with 5 mL of cold 6.1 N trichloroacetic acid (TCA), and incubated at 4°C overnight for protein precipitation. Tubes were then centrifuged at 3000 RCF for 30 minutes, supernatants discarded, and protein pellets washed twice with 90% acetone saturated with Tris base and once with 100% acetone. Pellets were air-dried, resuspended in 0.5 mL of 2X Laemmli Sample Buffer (Bio Rad), and incubated at 95°C for 10 minutes. Samples were then cooled to room temperature and centrifuged at 21,130 RCF for 10 minutes to remove remaining insoluble material for better gel resolution. 10 µL of each sample was loaded into the lanes of a Bio Rad Any kD SDS-PAGE gel. Gels were washed and stained with Coomassie Brilliant Blue R-250 and destained with water/methanol/acetic acid 50:40:10 until protein bands were clearly distinguishable from background. Gels was photographed with a Nikon Z30 on white background. Photographs were adjusted using Adobe Photoshop 2022 for exposure, levels, hue, saturation and/or contrast until fainter bands, easily visible by eye, were also distinguishable from background in the captured image. These adjustments were applied to the entire photograph equally.

### Semi-quantitative *In vitro* measurement of *E. faecalis* GAGase

A recently-published semi-quantitative turbidimetric assay of GAGase activity was used to examine the profiles of wild-type *E. faecalis* V587, Δ*hylA*Δ*hylB,* and Δ*hylA*Δ*hylB* with each constitutive expression construct (29). *E. faecalis* strains for GAG degradation assays were cultured overnight at 37°C in 3 mL of BHI with 20 µg/mL of chloramphenicol for plasmid containing strains. Cultures were then diluted 1:4 in BHI, adjusted to an OD_600_ nm of 0.04, pelleted at 6010 RCF for 8 minutes, and the supernatant was removed. Cell pellets were resuspended in 1 mL of 0.5X TSB containing 2.5 mg/mL of either hyaluronic acid (Sigma-Aldrich, catalog # 53747), chondroitin sulfate A (Sigma-Aldrich, catalog # C9819), or heparin sodium salt (Fisher BioReagents, catalog # BP2425), and 300 µL aliquoted in triplicate into 96-well plates. Plates were incubated in a hypoxia chamber for 24-hours at 35°C with 10% CO_2_ and 5% O_2_, followed by a semi-quantitative GAG degradation measurement as previously described (29). Briefly, the OD_600_ nm of all replicates was measured, bacteria were pelleted by centrifugation at 3214 RCF for 10 minutes, 180 µL of supernatant was transferred from each well to a new 96-well plate, and supernatants were then serially diluted in 1X PBS. 10 µL of 10% bovine serum albumin (BSA) was then added to all wells and OD_600_ was measured. Finally, 40 µL of 2M acetic acid was added to all wells, mixed by gentle stirring with the pipette tip, and OD_600_ was measured again. The percentage of GAG remaining was calculated by comparing the post-acetic acid OD_600_ values of all experimental conditions to bacteria-free controls (29). Percentage GAG remaining was calculated using the following formula using the OD_600_ reading of the post-acetic acid serial dilution falling in the center of the linear range of the assay for each condition.

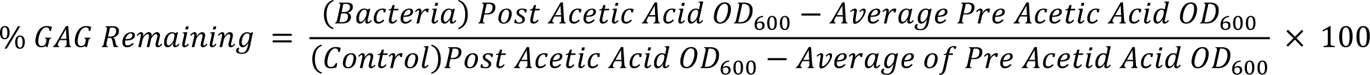

### Agarose gel electrophoresis for determination of GAG degradation product size

100 µL of indicated *E. faecalis* overnights were diluted into 20 mL of BHI containing appropriate antibiotics and 1 mg/mL HA (Sigma-Aldrich), incubated for 24 hours at 37°C with shaking, pelleted, and supernatants were filter sterilized. Two uninoculated HA BHI tubes with 20 µg/mL chloramphenicol were incubated the same way was included as a negative control, one of which had no pH adjustment, and one was acidified to pH 5.7 using 85-90% lactic acid (BeanTown Chemical) to simulate *E. faecalis* fermentation. Hyaluronidase positive controls were made by incubating HA BHI with either 12.5 µg/mL Bovine Hyaluronidase (Sigma-Aldrich) or *Streptomyces hyalurolyticus* hyaluronidase (Sigma-Aldrich) at 3 units/mL. Agarose gel electrophoresis was followed using a modified protocol from Echelon Biosciences (79) to determine HA size ranges. 750 µL of each supernatant was added to 250 µL of loading buffer (0.02% Bromophenol blue, 2 M sucrose in 1X Tris Borate EDTA (TBE) buffer), 10 µL of which was added to each well of a 2% (w/v) agarose gel and electrophoresis was performed at 100 V in TBE buffer. Gels were washed with 30% (v/v) ethanol for ∼15 minutes to fix the HA fragments within the gel prior to staining, and then washed in 50% (v/v) ethanol for ∼1 hour to remove the aqueous buffer as Stains-All dye has low water solubility. Gels were transferred to 0.01% Stains-All in 50% ethanol (Alfa Aesar) overnight, then destained with 50% ethanol until bands were clearly distinguishable from background staining. Gels were imaged on a ChemiDoc MP (BioRad) using the DyLight 680 setting.

The gel assay was also utilized to determine what HA size ranges were produced by recombinant HylB. For these experiments, 1 mg/ml of 601 kDa HA (Echelon Biosciences, catalog # HYA-601KEF-1) was incubated with either 27.5 µg/mL of recombinant HylB, 3 units/mL of *Streptomyces* hyaluronidase, or 50 µg/mL *E. coli* protein overnight at 37°C, and gels were prepared as above.

### Production of *E. faecalis* concentrated cell free supernatants for determination of GAG degradation by secreted products

To collect *E. faecalis* protein supernatants, 50 ml conical tubes containing 20 mL of BHI were inoculated with 100 µL of Δ*hylA*Δ*hylB* with pAOJ86, pAOJ84, or pAOJ55 and incubated for 4 hours with shaking at 37°C. After incubation, cultures were centrifuged and supernatants were filter sterilized and concentrated using a 10 kDa molecular weight cutoff filter. Retentates were then mixed with an equal proportion of 2 mg/mL HA in 20 mM sodium phosphate (pH 7) with 77 mM sodium chloride. A blank containing only the HA in phosphate buffer mixed 1:1 with PBS was also used. All samples were incubated for 24 hours statically at 37°C then subjected to agarose gel electrophoresis as described above.

### Cloning, expression, and purification of HylB Myc-6x His

The HylB expression vector pAOJ66 was generated by amplifying the *ef0818* gene with primers AOJ_651 and AOJ_652 and cloning into NcoI-HF/ApaI digested pBAD/myc his A (Invitrogen). The vector was propagated in *E. coli* TOP10, then transformed into *E. coli* BL21(DE3) pLysS (80) and cultured overnight in 50 mL LB with 100 µg/mL carbenicillin (Research Products International catalog # C46000) and 20 µg/mL chloramphenicol. The following day, 10 mL aliquots were sub-cultured into four 2 L flasks containing 900 mL pre-warmed Terrific Broth (81) supplemented with 10 mM of MgCl_2_ and 100 µg/mL carbenicillin. Flasks were incubated with shaking at 37°C until the OD_600_ reached 0.4-0.5 (∼3 hours post inoculation), after which 100 mL of 200 mM arabinose was added to each flask (20 mM final concentration) to induce protein expression (82). Cultures were incubated for 4 hours at 37°C with shaking, centrifuged at 8000 RCF for 10 minutes, and pellets were stored at -80°C overnight.

Cell pellets were thawed on ice, suspended in 200 mL total volume of lysis buffer (PBS pH 7.4 with 35 µM polymyxin B, 2% Triton X-100, 2 SIGMAFAST EDTA-free protease inhibitor tablets (Sigma-Aldrich, catalog # S8830-20TAB), 20 µL Benzonase nuclease (Sigma-Aldrich), and 50 mg of egg white lysozyme (Gold Biotechnology)), then incubated at 37°C for 30 with gentle agitation to lyse the cells. Lysates were centrifuged at 8000 RCF at 4°C for 30 minutes to remove insoluble material. The supernatant was filtered through a 0.22 µm PES membrane, and mixed with cold ammonium sulfate solution in PBS (pH 6) to a final concentration of 20% (v/v) ammonium sulfate. Samples were then centrifuged at 14500 RCF for 45 minutes at 4°C to remove higher molecular weight contaminants, supernatants were mixed with ammonium sulfate solution to a final concentration of 75% (v/v), and protein was precipitated overnight at 4°C. The next day, supernatants were centrifuged for 30 minutes at 14500 RCF at 4°C, the resulting protein pellet was dissolved in 100 mL of cold PBS pH 7.4 with 10 mM imidazole, and passed through a 3 mL His-Pur Ni-NTA column (Thermo Scientific, catalog # 88226) according to manufacturer instruction. The column-bound protein was eluted with 10 mL of 250 mM of imidazole in PBS pH 7.4, washed with PBS on a 10 kDa molecular weight cutoff column to remove imidazole, further washed with PBS + 5% glycerol, and the retentate was snap frozen and stored at -80°C.

Concentration and purity of recombinant protein was estimated using the Blue Dry Western technique (83) via a Trans-Blot Turbo RTA Transfer Kit, PVDF (Bio Rad, catalog # 1704272), and Trans-Blot Turbo transfer apparatus (Bio Rad) on the “large protein” pre-programmed setting. The blot was stained with the primary mouse α-c-Myc Monoclonal 9E10 antibody (Invitrogen, catalog # 13-2500) and Goat anti-mouse IgG1 Cross-Adsorbed Horseradish Peroxidase secondary antibody (Invitrogen, catalog # A10551), imaged colorimetrically (for Coomassie Brilliant Blue R-250), treated with Clarity Western ECL Substrate (Bio Rad, catalog # 170-5060), and imaged for chemiluminescence on a Bio Rad Chemi Doc MP imager. The chemiluminescence and colorimetric images were overlaid with the multi-image setting on the Chemi Doc MP to find bands contained the c-Myc tag of the recombinant protein. Densitometry was performed using Adobe Photoshop on raw colorimetric images to determine protein concentration by comparison to a Bovine Serum Albumin (BSA) dilution series of known concentrations loaded on the same gel.

### Assaying hyaluronidase activity of HylB Myc-6x His

A turbidimetric assay similar to the protocol developed by Sigma-Aldrich (84) was used to measure HA digestion by recombinant HylB. The assay consists of four reagents: Reagent A, the phosphate buffer solution, consisted of 20 mM sodium phosphate (pH 7) with 77 mM sodium chloride; Reagent B, the substrate solution, was HA sodium salt (Sigma-Aldrich) dissolved to a concentration of 0.3 mg/mL in reagent A; Reagent C, the enzyme diluent, consisted of Reagent A with 0.01% (w/v) BSA; and Reagent D, an acidic stop solution, was made by dissolving 0.1% (w/v) BSA into a buffer consisting of 24 mM sodium acetate and 79 mM glacial acetic acid adjusted to a pH of 3.75 with 5 M hydrochloric acid. Bovine Hyaluronidase Type I-S (Sigma-Aldrich, catalog # H3506-500MG) was utilized for comparison as a positive control, and was diluted to a concentration of 12.5 µg/mL in Reagent C. HA incubated with BSA at 15 µg/mL was used as the negative control for calculation of HA degradation, and soluble *E. coli* protein lysate from BL21(DE3) pLysS (pBAD-PL-Lux) at 50 µg/mL in Reagent C was included as an additional control. Protein samples were mixed with an equal volume of Reagent B and incubated at 37°C for 15 minutes, then the reaction was stopped by adding an equal volume of Reagent D. Remaining high molecular weight HA was precipitated during a 10 minute incubation at ambient temperature, the OD_600_ nm was read on a Synergy H1 plate reader (BioTek), and the percentage of remaining HA was estimated by dividing the OD_600_ nm of the BSA sample and multiplying by 100.

### *E. faecalis* growth curves

Growth in rich medium of WT V597, Δ*hylA,* Δ*hylB*, Δ*hylA*Δ*hylB,* or Δ*hylA*Δ*hylB* (pAOJ84) was measured by diluting overnight cultures 1:100 in BHI, TSB, or 0.5X TSB, aliquoting into a 96 well flat bottom plate, and incubating for 18-24 hours at 37°C with double-orbital shaking in a Synergy H1 plate reader (BioTek). Where indicated, media were supplemented with 2.5 mg/mL unfractionated HA or 2.5 mg/mL ∼5 kDa HA. Cell density was measured at 600 nm every 15 minutes.

To determine if *E. faecalis* V587 could use HA as a sole source of carbon, the Simplified Defined Media (SDM) recipe was used (47). Overnight cultures of *E. faecalis* V587 were centrifuged, washed once with an equal volume of PBS to remove traces of BHI, and diluted 1:100 in SDM with one of the following carbohydrate sources: 12 mM glucose (21), a two fold dilution series of HA (Sigma-Aldrich) starting at 2.5 mg/mL, or water (no carbon source). Growth was assessed as above.

Growth in human urine was assessed as previously described (40). Briefly, overnight cultures of *E. faecalis* were washed with PBS and diluted to an approximate cell density of 2 x 10^7^ CFU/mL in sterile, pooled urine from healthy female donors (Lee BioSolutions catalog # 991-03-P-FTD, lot # 01J5567) and incubated at 37°C with shaking. CFU/mL was determined by dilution and plating every hour for 7 hours.

### RAW-Blue NF-κB reporter assay

NF-κB activation by *E. faecalis* was assessed using RAW-Blue cells as previously described (50). RAW-Blue cells were obtained from InvivoGen (catalog # raw-sp) and routinely cultured at 37°C in a 5% CO_2_ in pre-warmed Dulbecco’s Modified Eagle’s Medium (DMEM, Corning catalog # 10-017-CM) with 4.5 g/L glucose, 0.584 g/L L-glutamine, 10% fetal bovine serum (Sigma-Aldrich), and no pyruvate. For each experiment, low-passage vials of cells were thawed and incubated in Corning Falcon 75 cm^2^ vented culture flasks at a density of 2 x 10^6^ / 15 mL of DMEM with 100 µg/mL Normocin (InvivoGen catalog # ant-nr) for approximately 72 hours. Culture media was then removed, cells were dislodged with a cell scraper and suspended in 10 mL of DMEM with 200 µg/mL Zeocin (InvivoGen catalog # ant-zn), counted on a Beckman Coulter Z2 Particle Counter and Size Analyzer set to a 10-20 µm size range, diluted to 5 x 10^5^ cells/mL in DMEM with 200 µg/mL Zeocin and 200 µL (1 x 10^5^ cells), and 100 µl were seeded into a Corning Falcon 96 well flat bottom polystyrene tissue culture treated plate and incubated overnight. The following day, the media was removed, the RAW-Blue cells were washed once with Dulbecco’s Phosphate Buffered Saline (DPBS) and then incubated with either LPS and/or HA disaccharide sodium salt (Biosynth catalog # OH11537) or infected with *E. faecalis* at a multiplicity of infection (MOI) of 10 (1 x 10^6^ CFU/well) in either DMEM or DMEM plus additives. MOI 10 was chosen as values higher than 10 were observed to acidify the media in a dose-dependent manner, low extracellular pH has been shown by to interfere with the NF-κB pathway (52–54). For infection experiments, media conditions used were: DMEM, DMEM with 100 µg/mL HA sodium salt at a molecular weight range of 500-749 kilodaltons (Lifecore Biomedical, catalog # HA700K), DMEM with 10 ng/mL ultrapure lipopolysaccharide (LPS) from *E. coli* O111:B4 (InvivoGen, catalog # tlrl-3pelps), or DMEM with both HA and LPS.

For all experiments, 200 µL of the treatment media or inoculum was added to the washed RAW-Blue cells and incubated for 6 hours. The plate was then centrifuged at 3000 RCF for 10 minutes, and 20 µL of supernatant was added to 180 µL of QUANTI-Blue Alkaline Phosphatase Detection Medium (InvivoGen catalog # rep-qbs), incubated at 37°C with shaking, and OD_640_ was read every 15 minutes for up to 16 hours to identify the optimal linear range. Under our experimental conditions, a 2 hour incubation was found to produce reads near the end of the linear range of the reporter assay, so this time point was used for reporting NF-κB activation.

### Analysis of cytokines, chemokines and growth factors from mouse bladders and kidneys

Flash frozen tissue homogenates were thawed on ice and pooled by infection date and inoculation group (WT, Δ*hylA*Δ*hylB*, or mock infected with PBS). Pooled samples were extracted using the Thermo Scientific Tissue Protein Extraction (T-PER) reagent bundle (Fisher Scientific catalog # B90002). 600 µL of total pooled sample volume was lysed with 700 µL T-PER lysis buffer with 1 Pierce Protease Inhibitor Mini Tablet per 7 mL of T-PER lysis buffer. Vortexed samples were centrifuged for 5 minutes at 10,000 RCF at 4°C and the supernatant removed. The protein concentration of 20 µL of supernatant was determined with Pierce Rapid Gold BCA Kit. Individual pooled samples were diluted to obtain 10 µg total protein in 50% T-PER buffer, 50% PBS.

Quantification of 32 different analytes in the adjusted bladder and kidney pools was provided by the Flow and Immune Analysis Shared Resource at the Roswell Park Comprehensive Cancer Center. The following analytes were measured: Eotaxin, G-CSF, GM-CSF, IFNγ, IL-1α, IL-1β, IL-2, IL-3, IL-4, IL-5, IL-6, IL-7, IL-9, IL-10, IL-12 (p40), IL-12 (p70), IL-13, IL-15, IL-17, IP-10, KC, LIF, LIX, MCP-1, M-CSF, MIG, MIP-1α, MIP-1β, MIP-2, RANTES, TNFα, and VEGF, using a commercially available kit (MCYTMAG-70K-PX32, Millipore Sigma, Burlington, Massachusetts). Data was acquired on a Luminex 200 instrument (Luminex Corporation, Austin, Texas). The experiment and instrument set-up were performed based on the manufacturer’s kit instructions. In brief, serially diluted standards were analyzed in duplicate wells, while the experimental samples were tested in single wells. The plate was incubated overnight with the multiplex beads on a plate shaker, in the dark, at 4°C and processed with the reporter reagents the next day, as per manufacturer’s instructions. Analyte concentrations were determined by extrapolating individual experimental fluorescence intensity values against each analyte’s standard curve using the BeadView multiplex data analysis software, version 1.0, (Upstate Cell Signaling Solutions, Lake Placid, New York). The analytical performance was checked using high concentration and low concentration quality controls provided with the kit of which the determined concentration for each analyte needs to be within the manufacturer-determined concentration range.

Radar charts were developed for select Luminex analytes to visualize the multivariate and multi-scale data simultaneously on a single plot. Separate plots were made for 6 hours and 24 hours using the same grid. The upper bound of the range (outer ring) for each variable was set as the log of the maximum average pg/ml values across replicates for V587, Δ*hylA* Δ*hylB* and PBS. The inner rings represent 25%, 50%, and 75% of the maximum, respectively Radar charts were developed in the R programming language using the “fmsb” packagev (85, 86).

### IL-6 ELISA

A colorimetric Mouse IL6 ELISA kit (Novus Biologicals NBP1-92668) was used to measure IL-6 from mouse tissue homogenates following the manufacturer instructions. Briefly, the ELISA plate was washed, bladder homogenates were thawed on ice and gently vortexed, and 50 µl of homogenate was mixed with 50 µl of diluent. 50 µl of Biotin conjugate was added to all wells (samples, blanks, and standards), the plate was incubated at room temperature for two hours with shaking at 400 RPM, then washed 5 time with a multichannel pipette. Strep-HRP was added and incubated for 1 hour at room temperature with shaking at 400 RPM. The plate was then washed, developer was added, and OD_620_ was monitored every minute on a BioTeck Synergy H1 until Standard 1 reached OD 0.9. Stop solution was then added and absorbance was read at 450 nm.

### Statistical analysis

Statistical analysis was performed using GraphPad Prism Software version 9. Significance was determined using a *P* < 0.05. All *P* values are two tailed at a 95% confidence interval. For colonization data in both the CAUTI and bacteremia model, we performed two types of statistical analysis to assess fitness of wild-type versus mutant strains. Data were first analyzed by Mann Whitney U-Test (*P*_MW_ < 0.05). Bacterial infection data does not necessarily follow distributions that are simple to perform standard statistical analysis on (87). To mitigate this issue, in addition to these tests, contingency tables of values above and below certain thresholds were constructed and analyzed for statistical significance by Fisher’s exact test (*P*_Fe_ < 0.05). In the CAUTI model a threshold of 1 x 10^5^ CFU per gram of tissue was chosen for the bladders / kidneys because it represented a middle point between the limit of detection (10^2^ CFU) and the upper range of recovered CFU (10^8^). The exception to this was at 6 HPI, in which the threshold was set to 1 x 10^4^ CFU due to lower overall colonization at this time point. For incidence of bacterial dissemination to the blood from CAUTI, the threshold was set at the limit of detection in the spleen (200 CFUs). For the bacteremia model in which mice were inoculated via tail-vein injection, statistical analysis was performed as in the CAUTI samples, with the exception that a cutoff of 1 x 10^5^ CFU per gram of tissue was used for all organs.

Statistical analysis of relative expression was performed with Wilcoxon signed rank test against a hypothetical value of 1. Analysis of growth curves, the NF-κB reporter assay with diHA, and the Luminex assay was performed using a two-way ANOVA for overall trends, with Dunnett’s Multiple Comparisons used to compare specific strains/conditions. The semi-quantitative GAG assays were analyzed with one-way ANOVA with Dunnett’s Multiple Comparisons.

## Acknowledgements

We would like to thank Dr. Thomas Russo and members of his laboratory for helpful comments and critiques, as well as Dr. Elsa Bou Ghanem for her useful suggestions. This work was funded by the National Institute of Diabetes and Digestive and Kidney Diseases under award R01 DK123158 to CEA, by university start-up funds to CEA, and by the Welch Foundation, award number AT-2030-20200401 to ND. Cytokine and chemokine analysis was performed by the Flow and Immune Analysis Shared Resource at the Roswell Park Comprehensive Cancer Center (Supported in part by NCI Cancer Center Support Grant 5P30 CA016056 and NCI R50 R50CA211108). The content is solely the responsibility of the authors and does not necessarily represent the official views of the National Institutes of Health.

## References

1. Weiner-Lastinger LM, Abner S, Benin AL, Edwards JR, Kallen AJ, Karlsson M, Magill SS, Pollock D, See I, Soe MM, Walters MS, Dudeck MA. 2020. Antimicrobial-resistant pathogens associated with pediatric healthcare-associated infections: Summary of data reported to the National Healthcare Safety Network, 2015-2017. Infect Control Hosp Epidemiol 41:19–30.

2. Weiner-Lastinger LM, Abner S, Edwards JR, Kallen AJ, Karlsson M, Magill SS, Pollock D, See I, Soe MM, Walters MS, Dudeck MA. 2020. Antimicrobial-resistant pathogens associated with adult healthcare-associated infections: Summary of data reported to the National Healthcare Safety Network, 2015-2017. Infect Control Hosp Epidemiol 41:1–18.

3. Armbruster CE, Prenovost K, Mobley HL, Mody L. 2017. How Often Do Clinically Diagnosed Catheter-Associated Urinary Tract Infections in Nursing Homes Meet Standardized Criteria? J Am Geriatr Soc 65:395–401.

4. Werneburg GT. 2022. Catheter-Associated Urinary Tract Infections: Current Challenges and Future Prospects. Res Rep Urol 14:109–133.

5. Klevens RM, Edwards JR, Richards CL, Jr., Horan TC, Gaynes RP, Pollock DA, Cardo DM. 2007. Estimating health care-associated infections and deaths in U.S. hospitals, 2002. Public Health Rep 122:160–6.

6. Basu A, Patel NG, Nicholson ED, Weiss RJ. 2022. Spatiotemporal diversity and regulation of glycosaminoglycans in cell homeostasis and human disease. Am J Physiol Cell Physiol 322:C849–C864.

7. Rügheimer L, Olerud J, Johnsson C, Takahashi T, Shimizu K, Hansell P. 2009. Hyaluronan synthases and hyaluronidases in the kidney during changes in hydration status. Matrix Biology 28:390–395.

8. Toole BP. 2000. Hyaluronan is not just a goo! J Clin Invest 106:335–6.

9. Karamanos NK, Theocharis AD, Piperigkou Z, Manou D, Passi A, Skandalis SS, Vynios DH, Orian-Rousseau V, Ricard-Blum S, Schmelzer CEH, Duca L, Durbeej M, Afratis NA, Troeberg L, Franchi M, Masola V, Onisto M. 2021. A guide to the composition and functions of the extracellular matrix. Febs j 288:6850–6912.

10. Jiang D, Liang J, Noble PW. 2007. Hyaluronan in tissue injury and repair. Annu Rev Cell Dev Biol 23:435–61.

11. Fan J, Sun Y. 2019. Ultra-Structure of Endothelial Surface Glycocalyx Revealed by Stochastic Optical Reconstruction Microscopy (STORM). Biorheology 56:77–88.

12. Reitsma S, Slaaf DW, Vink H, van Zandvoort MA, oude Egbrink MG. 2007. The endothelial glycocalyx: composition, functions, and visualization. Pflugers Arch 454:345–59.

13. Klingler CH. 2016. Glycosaminoglycans: how much do we know about their role in the bladder? Urologia 83 Suppl 1:11–4.

14. Parsons CL, Mulholland SG, Anwar H. 1979. Antibacterial activity of bladder surface mucin duplicated by exogenous glycosaminoglycan (heparin). Infect Immun 24:552–7.

15. Parsons CL, Greenspan C, Moore SW, Mulholland SG. 1977. Role of surface mucin in primary antibacterial defense of bladder. Urology 9:48–52.

16. Ruggieri MR, Hanno PM, Levin RM. 1984. The effects of heparin on the adherence of five species of urinary tract pathogens to urinary bladder mucosa. Urol Res 12:199–203.

17. Jin C, Zong Y. 2023. The role of hyaluronan in renal cell carcinoma. Front Immunol 14:1127828.

18. Kolar SL, Kyme P, Tseng CW, Soliman A, Kaplan A, Liang J, Nizet V, Jiang D, Murali R, Arditi M, Underhill DM, Liu GY. 2015. Group B Streptococcus Evades Host Immunity by Degrading Hyaluronan. Cell Host Microbe 18:694–704.

19. Wang Z, Guo C, Xu Y, Liu G, Lu C, Liu Y. 2014. Two novel functions of hyaluronidase from Streptococcus agalactiae are enhanced intracellular survival and inhibition of proinflammatory cytokine expression. Infect Immun 82:2615–25.

20. Hynes WL, Walton SL. 2000. Hyaluronidases of Gram-positive bacteria. FEMS Microbiol Lett 183:201–7.

21. Marion C, Stewart JM, Tazi MF, Burnaugh AM, Linke CM, Woodiga SA, King SJ. 2012. Streptococcus pneumoniae can utilize multiple sources of hyaluronic acid for growth. Infect Immun 80:1390–8.

22. Rosan B, Williams NB. 1964. Hyaluronidase Production by Oral Enterococci. Arch Oral Biol 9:291–8.

23. Sherman JM. 1937. The Streptococci. Bacteriol Rev 1:3–97.

24. Lebreton F WR, Gilmore MS. 2014. Enterococcus Diversity, Origins in Nature, and Gut Colonization. . In Gilmore MS CD, Ike Y, et al., editors. (ed), Enterococci: From Commensals to Leading Causes of Drug Resistant Infection Massachusetts Eye and Ear Infirmary, Internet.

25. Rice LB, Carias L, Rudin S, Vael C, Goossens H, Konstabel C, Klare I, Nallapareddy SR, Huang W, Murray BE. 2003. A potential virulence gene, hylEfm, predominates in Enterococcus faecium of clinical origin. J Infect Dis 187:508–12.

26. Arias CA, Panesso D, Singh KV, Rice LB, Murray BE. 2009. Cotransfer of antibiotic resistance genes and a hylEfm-containing virulence plasmid in Enterococcus faecium. Antimicrob Agents Chemother 53:4240–6.

27. Laverde Gomez JA, van Schaik W, Freitas AR, Coque TM, Weaver KE, Francia MV, Witte W, Werner G. 2011. A multiresistance megaplasmid pLG1 bearing a hylEfm genomic island in hospital Enterococcus faecium isolates. Int J Med Microbiol 301:165–75.

28. Panesso D, Montealegre MC, Rincon S, Mojica MF, Rice LB, Singh KV, Murray BE, Arias CA. 2011. The hylEfm gene in pHylEfm of Enterococcus faecium is not required in pathogenesis of murine peritonitis. BMC Microbiol 11:20.

29. Nguyen VH, Khan F, Shipman BM, Neugent ML, Hulyalkar NV, Cha NY, Zimmern PE, De Nisco NJ. 2021. A Semi-Quantitative Assay to Measure Glycosaminoglycan Degradation by the Urinary Microbiota. Front Cell Infect Microbiol 11:803409.

30. Olson RD, Assaf R, Brettin T, Conrad N, Cucinell C, Davis JJ, Dempsey DM, Dickerman A, Dietrich EM, Kenyon RW, Kuscuoglu M, Lefkowitz EJ, Lu J, Machi D, Macken C, Mao C, Niewiadomska A, Nguyen M, Olsen GJ, Overbeek JC, Parrello B, Parrello V, Porter JS, Pusch GD, Shukla M, Singh I, Stewart L, Tan G, Thomas C, VanOeffelen M, Vonstein V, Wallace ZS, Warren AS, Wattam AR, Xia F, Yoo H, Zhang Y, Zmasek CM, Scheuermann RH, Stevens RL. 2022. Introducing the Bacterial and Viral Bioinformatics Resource Center (BV-BRC): a resource combining PATRIC, IRD and ViPR. Nucleic Acids Res doi:10.1093/nar/gkac1003.

31. Sahm DF, Kissinger J, Gilmore MS, Murray PR, Mulder R, Solliday J, Clarke B. 1989. In vitro susceptibility studies of vancomycin-resistant *Enterococcus faecalis*. Antimicrob Agents Chemother 33:1588–91.

32. Tatusova T, DiCuccio M, Badretdin A, Chetvernin V, Nawrocki EP, Zaslavsky L, Lomsadze A, Pruitt KD, Borodovsky M, Ostell J. 2016. NCBI prokaryotic genome annotation pipeline. Nucleic Acids Res 44:6614–24.

33. Olson RD, Assaf R, Brettin T, Conrad N, Cucinell C, Davis JJ, Dempsey DM, Dickerman A, Dietrich EM, Kenyon RW, Kuscuoglu M, Lefkowitz EJ, Lu J, Machi D, Macken C, Mao C, Niewiadomska A, Nguyen M, Olsen GJ, Overbeek JC, Parrello B, Parrello V, Porter JS, Pusch GD, Shukla M, Singh I, Stewart L, Tan G, Thomas C, VanOeffelen M, Vonstein V, Wallace ZS, Warren AS, Wattam AR, Xia F, Yoo H, Zhang Y, Zmasek CM, Scheuermann RH, Stevens RL. 2023. Introducing the Bacterial and Viral Bioinformatics Resource Center (BV-BRC): a resource combining PATRIC, IRD and ViPR. Nucleic Acids Res 51:D678–d689.

34. Almagro Armenteros JJ, Tsirigos KD, Sonderby CK, Petersen TN, Winther O, Brunak S, von Heijne G, Nielsen H. 2019. SignalP 5.0 improves signal peptide predictions using deep neural networks. Nat Biotechnol 37:420–423.

35. Navarre WW, Schneewind O. 1994. Proteolytic cleavage and cell wall anchoring at the LPXTG motif of surface proteins in gram-positive bacteria. Mol Microbiol 14:115–21.

36. Lu S, Wang J, Chitsaz F, Derbyshire MK, Geer RC, Gonzales NR, Gwadz M, Hurwitz DI, Marchler GH, Song JS, Thanki N, Yamashita RA, Yang M, Zhang D, Zheng C, Lanczycki CJ, Marchler-Bauer A. 2020. CDD/SPARCLE: the conserved domain database in 2020. Nucleic Acids Res 48:D265–D268.

37. Jumper J, Evans R, Pritzel A, Green T, Figurnov M, Ronneberger O, Tunyasuvunakool K, Bates R, Žídek A, Potapenko A, Bridgland A, Meyer C, Kohl SAA, Ballard AJ, Cowie A, Romera-Paredes B, Nikolov S, Jain R, Adler J, Back T, Petersen S, Reiman D, Clancy E, Zielinski M, Steinegger M, Pacholska M, Berghammer T, Bodenstein S, Silver D, Vinyals O, Senior AW, Kavukcuoglu K, Kohli P, Hassabis D. 2021. Highly accurate protein structure prediction with AlphaFold. Nature 596:583–589.

38. Varadi M, Anyango S, Deshpande M, Nair S, Natassia C, Yordanova G, Yuan D, Stroe O, Wood G, Laydon A, Žídek A, Green T, Tunyasuvunakool K, Petersen S, Jumper J, Clancy E, Green R, Vora A, Lutfi M, Figurnov M, Cowie A, Hobbs N, Kohli P, Kleywegt G, Birney E, Hassabis D, Velankar S. 2021. AlphaFold Protein Structure Database: massively expanding the structural coverage of protein-sequence space with high-accuracy models. Nucleic Acids Research 50:D439–D444.

39. Innocenti N, Golumbeanu M, Fouquier d’Herouel A, Lacoux C, Bonnin RA, Kennedy SP, Wessner F, Serror P, Bouloc P, Repoila F, Aurell E. 2015. Whole-genome mapping of 5’ RNA ends in bacteria by tagged sequencing: a comprehensive view in Enterococcus faecalis. RNA 21:1018–30.

40. Learman BS, Brauer AL, Eaton KA, Armbruster CE. 2019. A Rare Opportunist, *Morganella morganii*, Decreases Severity of Polymicrobial Catheter-Associated Urinary Tract Infection. Infect Immun 88.

41. Honda T, Kaneiwa T, Mizumoto S, Sugahara K, Yamada S. 2012. Hyaluronidases Have Strong Hydrolytic Activity toward Chondroitin 4-Sulfate Comparable to that for Hyaluronan. Biomolecules 2:549–563.

42. Duan J, Kasper DL. 2010. Oxidative depolymerization of polysaccharides by reactive oxygen/nitrogen species. Glycobiology 21:401–409.

43. Scheibner KA, Lutz MA, Boodoo S, Fenton MJ, Powell JD, Horton MR. 2006. Hyaluronan fragments act as an endogenous danger signal by engaging TLR2. J Immunol 177:1272–81.

44. Termeer C, Benedix F, Sleeman J, Fieber C, Voith U, Ahrens T, Miyake K, Freudenberg M, Galanos C, Simon JC. 2002. Oligosaccharides of Hyaluronan activate dendritic cells via toll-like receptor 4. J Exp Med 195:99–111.

45. Pritchard DG, Lin B, Willingham TR, Baker JR. 1994. Characterization of the group B streptococcal hyaluronate lyase. Arch Biochem Biophys 315:431–7.

46. Ohya T, Kaneko Y. 1970. Novel hyaluronidase from streptomyces. Biochim Biophys Acta 198:607–9.

47. Khan H, Flint SH, Yu PL. 2013. Development of a chemically defined medium for the production of enterolysin A from Enterococcus faecalis B9510. J Appl Microbiol 114:1092–102.

48. Coleman M, Armistead B, Orvis A, Quach P, Brokaw A, Gendrin C, Sharma K, Ogle J, Merillat S, Dacanay M, Wu TY, Munson J, Baldessari A, Vornhagen J, Furuta A, Nguyen S, Adams Waldorf KM, Rajagopal L. 2021. Hyaluronidase Impairs Neutrophil Function and Promotes Group B Streptococcus Invasion and Preterm Labor in Nonhuman Primates. mBio 12.

49. Vornhagen J, Quach P, Boldenow E, Merillat S, Whidbey C, Ngo LY, Adams Waldorf KM, Rajagopal L. 2016. Bacterial Hyaluronidase Promotes Ascending GBS Infection and Preterm Birth. mBio 7:e00781–16.

50. Tien BYQ, Goh HMS, Chong KKL, Bhaduri-Tagore S, Holec S, Dress R, Ginhoux F, Ingersoll MA, Williams RBH, Kline KA. 2017. Enterococcus faecalis Promotes Innate Immune Suppression and Polymicrobial Catheter-Associated Urinary Tract Infection. Infect Immun 85.

51. Kao PHN, Kline KA. 2019. Dr. Jekyll and Mr. Hide: How Enterococcus faecalis Subverts the Host Immune Response to Cause Infection. J Mol Biol 431:2932–2945.

52. Gerry AB, Leake DS. 2014. Effect of low extracellular pH on NF-kappaB activation in macrophages. Atherosclerosis 233:537–544.

53. Brenda Yin Qi T. 2020. Mechanism of immune modulation in macrophages by Enterococcus faecalis doi:doi:10.32657/10356/138251 Nanyang Technological University.

54. Heming TA, Tuazon DM, Dave SK, Chopra AK, Peterson JW, Bidani A. 2001. Post-transcriptional effects of extracellular pH on tumour necrosis factor-alpha production in RAW 246.7 and J774 A.1 cells. Clin Sci (Lond) 100:259–66.

55. Steiner B, Romero-Steiner S, Cruce D, George R. 1997. Cloning and sequencing of the hyaluronate lyase gene from Propionibacterium acnes. Can J Microbiol 43:315–21.

56. Farrell AM, Taylor D, Holland KT. 1995. Cloning, nucleotide sequence determination and expression of the Staphylococcus aureus hyaluronate lyase gene. FEMS Microbiol Lett 130:81–5.

57. Li S, Jedrzejas MJ. 2001. Hyaluronan binding and degradation by Streptococcus agalactiae hyaluronate lyase. J Biol Chem 276:41407–16.

58. Li S, Kelly SJ, Lamani E, Ferraroni M, Jedrzejas MJ. 2000. Structural basis of hyaluronan degradation by Streptococcus pneumoniae hyaluronate lyase. Embo j 19:1228–40.

59. Wang X, Zhang S, Wu H, Li Y, Yu W, Han F. 2021. Expression and characterization of a thermotolerant and pH-stable hyaluronate lyase from Thermasporomyces composti DSM22891. Protein Expr Purif 182:105840.

60. Gase K, Ozegowski J, Malke H. 1998. The Streptococcus agalactiae hylB gene encoding hyaluronate lyase: completion of the sequence and expression analysis. Biochim Biophys Acta 1398:86–98.

61. Sheldon WL, Macauley MS, Taylor EJ, Robinson CE, Charnock SJ, Davies GJ, Vocadlo DJ, Black GW. 2006. Functional analysis of a group A streptococcal glycoside hydrolase Spy1600 from family 84 reveals it is a beta-N-acetylglucosaminidase and not a hyaluronidase. Biochem J 399:241–7.

62. Alhajjar N, Chatterjee A, Spencer BL, Burcham LR, Willett JLE, Dunny GM, Duerkop BA, Doran KS. 2020. Genome-Wide Mutagenesis Identifies Factors Involved in Enterococcus faecalis Vaginal Adherence and Persistence. Infect Immun 88.

63. Soussan D, Salze M, Ledormand P, Sauvageot N, Boukerb A, Lesouhaitier O, Fichant G, Rincé A, Quentin Y, Muller C. 2023. The NagY regulator: A member of the BglG/SacY antiterminator family conserved in Enterococcus faecalis and involved in virulence. Frontiers in Microbiology 13.

64. Tanaka T, Narazaki M, Kishimoto T. 2014. IL-6 in inflammation, immunity, and disease. Cold Spring Harb Perspect Biol 6:a016295.

65. Guiton PS, Hung CS, Hancock LE, Caparon MG, Hultgren SJ. 2010. Enterococcal biofilm formation and virulence in an optimized murine model of foreign body-associated urinary tract infections. Infect Immun 78:4166–75.

66. Guiton PS, Hannan TJ, Ford B, Caparon MG, Hultgren SJ. 2013. Enterococcus faecalis Overcomes Foreign Body-Mediated Inflammation To Establish Urinary Tract Infections. Infection and Immunity 81:329–339.

67. McBride SM, Fischetti VA, Leblanc DJ, Moellering RC, Jr., Gilmore MS. 2007. Genetic diversity among Enterococcus faecalis. PLoS One 2:e582.

68. Shankar N, Lockatell CV, Baghdayan AS, Drachenberg C, Gilmore MS, Johnson DE. 2001. Role of Enterococcus faecalis surface protein Esp in the pathogenesis of ascending urinary tract infection. Infect Immun 69:4366–72.

69. Monk IR, Shah IM, Xu M, Tan MW, Foster TJ. 2012. Transforming the untransformable: application of direct transformation to manipulate genetically Staphylococcus aureus and Staphylococcus epidermidis. mBio 3.

70. Maguin E, Duwat P, Hege T, Ehrlich D, Gruss A. 1992. New thermosensitive plasmid for gram-positive bacteria. J Bacteriol 174:5633–8.

71. Toledo-Arana A, Valle J, Solano C, Arrizubieta MJ, Cucarella C, Lamata M, Amorena B, Leiva J, Penades JR, Lasa I. 2001. The enterococcal surface protein, Esp, is involved in Enterococcus faecalis biofilm formation. Appl Environ Microbiol 67:4538–45.

72. Armbruster CE, Smith SN, Johnson AO, DeOrnellas V, Eaton KA, Yep A, Mody L, Wu W, Mobley HLT. 2017. The Pathogenic Potential of Proteus mirabilis Is Enhanced by Other Uropathogens during Polymicrobial Urinary Tract Infection. Infect Immun 85.

73. Gaston JR, Andersen MJ, Johnson AO, Bair KL, Sullivan CM, Guterman LB, White AN, Brauer AL, Learman BS, Flores-Mireles AL, Armbruster CE. 2020. Enterococcus faecalis Polymicrobial Interactions Facilitate Biofilm Formation, Antibiotic Recalcitrance, and Persistent Colonization of the Catheterized Urinary Tract. Pathogens 9.

74. Eckert C, Lecerf M, Dubost L, Arthur M, Mesnage S. 2006. Functional analysis of AtlA, the major N-acetylglucosaminidase of Enterococcus faecalis. J Bacteriol 188:8513–9.

75. Salamaga B, Prajsnar TK, Jareño-Martinez A, Willemse J, Bewley MA, Chau F, Ben Belkacem T, Meijer AH, Dockrell DH, Renshaw SA, Mesnage S. 2017. Bacterial size matters: Multiple mechanisms controlling septum cleavage and diplococcus formation are critical for the virulence of the opportunistic pathogen Enterococcus faecalis. PLoS Pathog 13:e1006526.

76. Lizier M, Sarra PG, Cauda R, Lucchini F. 2010. Comparison of expression vectors in Lactobacillus reuteri strains. FEMS Microbiol Lett 308:8–15.

77. Papagianni M, Avramidis N, Filioussis G. 2007. High efficiency electrotransformation of Lactococcus lactis spp. lactis cells pretreated with lithium acetate and dithiothreitol. BMC Biotechnol 7:15.

78. Qin X, Singh KV, Weinstock GM, Murray BE. 2001. Characterization of fsr, a regulator controlling expression of gelatinase and serine protease in Enterococcus faecalis OG1RF. J Bacteriol 183:3372–82.

79. Author U. 2019. Protocol for Hyaluronic Acid Gel Electrophoresis.

80. Studier FW, Moffatt BA. 1986. Use of bacteriophage T7 RNA polymerase to direct selective high-level expression of cloned genes. J Mol Biol 189:113–30.

81. Anonymous. 2015. Terrific Broth (TB) Medium. Cold Spring Harbor Protocols 2015:pdb.rec085894.

82. Guzman LM, Belin D, Carson MJ, Beckwith J. 1995. Tight regulation, modulation, and high-level expression by vectors containing the arabinose PBAD promoter. J Bacteriol 177:4121–30.

83. Naryzhny SN. 2009. Blue Dry Western: simple, economic, informative, and fast way of immunodetection. Anal Biochem 392:90–5.

84. Author U. 1994. Enzymatic Assay of HYALURONATE LYASE (EC 4.2.2.1).

85. Nakazawa M. 2019. Package “fmsb’. https://cran.r-project.org/web/packages/fmsb/index.html. Accessed

86. Saary MJ. 2008. Radar plots: a useful way for presenting multivariate health care data. J Clin Epidemiol 61:311–7.

87. Armbruster CE, Smith SN, Mody L, Mobley HLT. 2018. Urine Cytokine and Chemokine Levels Predict Urinary Tract Infection Severity Independent of Uropathogen, Urine Bacterial Burden, Host Genetics, and Host Age. Infection and Immunity 86:10.1128/iai.00327-18.

